# Accounting for taste: A multi-attribute neurocomputational model explains the neural dynamics of choices for self and others

**DOI:** 10.1101/329391

**Authors:** Alison Harris, John A. Clithero, Cendri A. Hutcherson

## Abstract

How do we make choices for others with different preferences from our own? Although neuroimaging studies implicate similar circuits in representing preferences for oneself and others, some models propose that additional corrective mechanisms come online when choices for others diverge from one’s own preferences. Here we used event-related potentials (ERP) in humans, in combination with computational modeling, to examine how social information is integrated in the time leading up to choices for oneself and others. Hungry male and female participants with unrestricted diets selected foods for themselves, a similar unrestricted eater, and a dissimilar, self-identified healthy eater. Across choices for both oneself and others, ERP value signals emerged within the same time window but differentially reflected taste and health attributes based on the recipient’s preferences. Choices for the dissimilar recipient were associated with earlier activity localized to brain regions implicated in social cognition including temporoparietal junction (TPJ). Finally, response-locked analysis revealed a late ERP component specific to choices for the similar recipient, localized to the parietal lobe, that appeared to reflect differences in the response threshold based on uncertainty. A multi-attribute computational model supported the link between specific ERP components and distinct model parameters, and was not significantly improved by adding time-dependent dual processes. Model simulations suggested that longer response times (RTs) previously associated with effortful correction may alternatively arise from higher choice uncertainty. Together these results provide a parsimonious neurocomputational mechanism for social decision-making, additionally explaining divergent patterns of choice and RT data in decisions for oneself and others.

**Significance Statement:** How do we choose for others, particularly when they have different preferences? Whereas some studies suggest that similar neural circuits underlie decision-making for oneself and others, others argue for additional, slower perspective-taking mechanisms. Combining event-related potentials with computational modeling, we found that integration of others’ preferences occurs over the same timescale as for oneself, while differentially tracking recipient-relevant attributes. Although choosing for others took longer and produced differences in late-emerging neural responses, computational modeling attributed these patterns to greater response caution rather than egocentric bias correction. Computational simulations also correctly predicted when and why choosing differently for others takes longer, suggesting that a model incorporating value integration and evidence accumulation can parsimoniously account for complex patterns in social decision-making.

---

Whether in parenting or politics, we are often required to make decisions for other individuals whose preferences may differ substantially from our own. A critical question concerns whether and when such choices require perspective-taking mechanisms that are slower, more effortful, or computationally costly (Epley et al., 2004). For example, studies of response time (RT) find that people take longer to choose for similar others when those choices diverge from what they choose for themselves (Tamir and Mitchell, 2013; Apps et al., 2016; Volz et al., 2017), perhaps because slower, more controlled processing is required to adjust an otherwise automatic egocentric bias. Overcoming egocentric choice biases has also been associated with increased activation in brain regions associated with social cognition, such as the ventromedial prefrontal cortex (VMPFC) and temporoparietal junction (TPJ) (Tamir and Mitchell, 2010; Silani et al., 2013).

Yet recent work using computational modeling suggests that slower RTs and activation in VMPFC and TPJ during some social decision-making tasks can result from a noisy value-integration mechanism, without requiring additional slow or effortful cognitive processes (Hutcherson et al., 2015; Krajbich et al., 2015; Berkman et al., 2017). Moreover, several functional magnetic resonance imaging (fMRI) studies have found that constructing values and choices for oneself and others recruit the same regions of VMPFC, even when preferences differ substantially (Nicolle et al., 2012; Janowski et al., 2013). Unfortunately, the low temporal resolution of fMRI obscures the dynamics of this process, and recent computational models of choice for others (Devaine and Daunizeau, 2017; Tarantola et al., 2017) have not explicitly examined when and why constructing values for others might require longer, more extensive processing compared to the self.

This paper aims to answer three related questions. First, are additional neural computations involved when choosing for others with similar or dissimilar preferences? Second, what are the temporal dynamics of those computations? Third, how might these computations explain when and why people show egocentric biases, or instead make different choices for themselves and others? To address these questions, we use a dietary choice task (Figure 1) in which participants with unrestricted diets made food choices for themselves and two partners: a *Similar* partner with unrestricted dietary preferences, and a *Dissimilar* partner who self-identified as a healthy eater. We employed a novel combination of event-related potentials (ERP), computational modeling, and simulation to identify the processes necessary to account for the full pattern of choices, RTs, and neural responses observed when choosing for oneself, similar, and dissimilar others.

**Figure 1.**
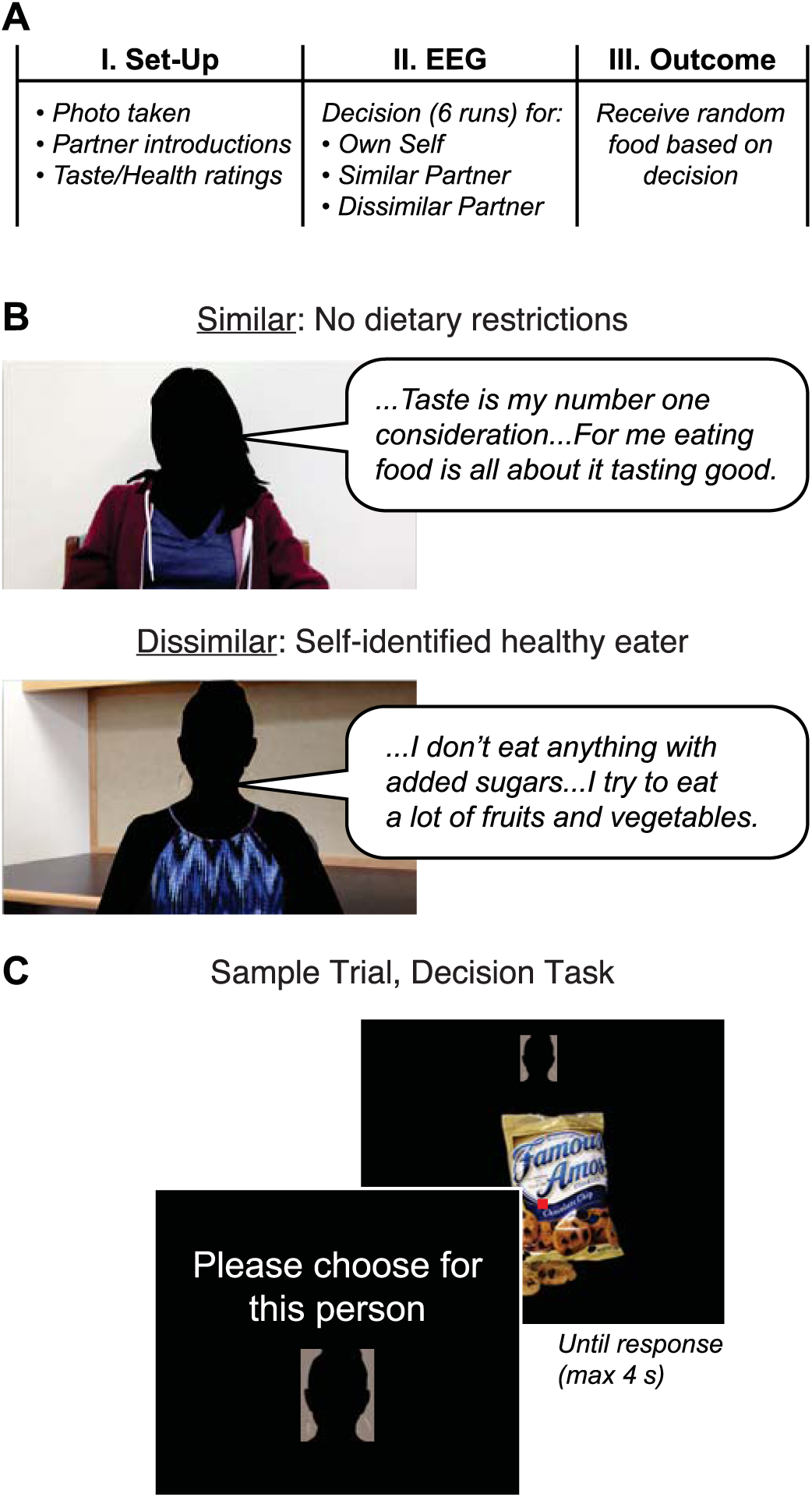
Social food choice task. (A) Experimental session. In Part 1, hungry participants with unrestricted diets were introduced to two partners, and provided taste and health ratings for the foods used in the experiment. Then, in Part 2, participants made food choices for themselves and the partners while their brain activity was measured with EEG. Participants knew that their choices mattered because a single trial was randomly selected and implemented for each recipient in Part 3. (B) Sample statements from the *Similar* (top) and *Dissimilar* (bottom) partners in the experiment. (C) Sample stimuli and screens from the decision task.

The best-fitting model, a multi-attribute version of the drift-diffusion model (DDM) (Smith and Ratcliff, 2004; Ratcliff and McKoon, 2008), suggests that individuals may indeed use their own preferences as a source of information, particularly when deciding for similar others. Yet a slow, serial adjustment mechanism is not required to explain when and why people choose differently for others and take longer to do so. Instead, an early-acting perspective-taking mechanism, localized to the temporoparietal junction (TPJ), may play a role in adjusting the importance of different stimulus attributes (e.g., tastiness and healthiness) based on inferences about the recipient’s preferences. These attributes are integrated into a noisy overall value signal, localized to a region of VMPFC previously implicated in value construction (Bartra et al., 2013; Clithero and Rangel, 2014) with a similar time course regardless of recipient. We find further evidence that these VMPFC value signals serve as input to an evidence accumulation process (Rangel and Clithero, 2013) that produces a choice when the accumulated evidence crosses a threshold whose height can be adjusted based on uncertainty about others’ preferences. Thus, although choices for other involve multiple distinct systems, these computations follow different temporal dynamics and play different functional roles than previously hypothesized. These results show that a parsimonious neurocomputational framework can account not only for when people choose differently for themselves than for others, but also when and why these choices are associated with additional processing time.

## Methods

### Participants

Thirty-eight right-handed participants (ages 18-40, 20 female) from the Claremont Colleges participated in exchange for course credit. Participants were screened to ensure that they enjoyed common appetitive snacks and did not self-identify as healthy eaters. Although all 38 participants were included in the behavioral model, two individuals were excluded from the combined behavioral-neurophysiological model due to excessively noisy EEG data in which cognitive signals could not be reliably separated from artefactual noise using artifact removal protocols (see “EEG data acquisition and preprocessing”). All participants provided written informed consent before participation. Procedures were reviewed and approved by the Claremont McKenna College Institutional Review Board.

### Stimuli

Stimuli consisted of color pictures of 250 common appetitive foods on a black background (Figure 1C; 576 × 432 pixels). Foods were chosen to span a full range of perceived tastiness and healthiness based on pilot ratings from a separate group of participants (N = 25). The stimulus set included fruits and vegetables, chips, and candy bars.

### Procedure

The experiment consisted of three parts (Figure 1A). In Part 1, participants received information about the two partners for whom they would be making food choices, after which they rated foods in terms of taste and health attributes. In Part 2, participants made choices about whether to eat these foods, separately for themselves and their two partners (*Own* self, *Similar* other, *Dissimilar* other conditions), while their brain activity was recorded with electroencephalography (EEG). Finally, in Part 3, a single trial was randomly selected for each recipient and the outcome of that trial was implemented.

Participants were instructed to fast for four hours prior to arrival to ensure they were hungry and motivated to perform the decision task. Compliance was assessed verbally before starting the experiment. Next, the experimenter took a photo of the participant against a white wall, which was employed as a visual cue for the trials in which the participant chose food for him/herself in Part 2. Participants then completed a short computer-based survey in which they indicated the extent to which they generally use health information in deciding what to eat, and made a series of twelve binary choices between a healthier and less healthy but tastier option (e.g., green salad vs. French fries).

Participants received instructions familiarizing them with the overall structure of the decision task and introducing the two partners (Figure 1B), matched to the participant’s own gender. Participants were informed that the partners were real people expressing their true dietary preferences, and that they had in fact agreed to eat at least three bites of whichever food the participant selected for them. This was true; no deception was used in the experiment. Each participant received information that one partner had unrestricted dietary preferences similar to their own (*Similar* partner; Figure 1B, top), while the other had quite different preferences (*Dissimilar* partner; Figure 1B, bottom). First, participants viewed two short video interviews in which each partner responded to prompts about their eating habits. Each partner was introduced by name via an intertitle at the beginning of the interview. During the interview, partners described their food preferences in general, and answered questions about specific choices (e.g., “Pizza or a salad”). Representative quotes for the partner without dietary restrictions included: “I would say for me, taste is my number one consideration when I’m eating…If it tastes good but it’s not healthy, I’ll just eat less of it, but for me eating food is all about it tasting good.” Representative quotes for the self-identified healthy eater included the following: “I don’t eat anything with added sugars…I try to eat a lot of fruits and vegetables, and I eat a lot of peanut butter and almond butter.” The video interviews were lightly edited for conciseness, but otherwise were not manipulated. To give participants a fuller picture of the partners’ preferences, they were also shown the partners’ responses to the same 12-item food questionnaire they had completed earlier.

Following introduction to the partners, participants completed two blocks of trials in which they rated each of 250 appetitive foods in terms of tastiness and healthiness. Each food was displayed until the participant entered a keypress response using a 4-point scale (1 = Strong No, 2 = Weak No, 3 = Weak Yes, 4 = Strong Yes). The order of the rating blocks and the left-to-right order of the response keys was counterbalanced across participants.

Next, EEG data were recorded while participants performed a simple decision-making task (Figure 1C) in which they chose whether or not a particular recipient (*Own* self, *Similar* other, or *Dissimilar* other) would have the opportunity to consume each food at the end of the experiment. Due to time constraints, this decision task used a subset of 200 foods (out of the original 250), chosen to span the full range of the participant’s taste and health ratings. Participants made decisions regarding each food both for themselves and the two partners, resulting in a total of 600 trials, in blocks of 10 trials per recipient. At the beginning of each block, the participant saw a prompt (“For the next set of trials, please choose for this person”) above a picture of the intended recipient. For *Own* self trials, this picture was the head shot taken upon the participant’s arrival at the lab; the other partners’ images were captured from screenshots of the video interviews. The duration of the recipient prompt was randomly jittered between 4 and 5 seconds. In addition, a smaller picture of the intended recipient was displayed at the top of the screen throughout the block. On each trial, participants saw a food image centered on the screen, which remained visible until a response was entered (maximum duration 4 s), using a 4-point scale (1 = Strong No, 2 = Weak No, 3 = Weak Yes, 4 = Strong Yes). This response method captures both the strength and direction of participants’ preferences (Hare et al., 2009; Hutcherson et al., 2015). The order of the recipient blocks and of the food images was randomized across participants. The task was subdivided into 6 runs, and within each run the blocks of trials were further subdivided by intervening self-paced breaks. During the decision trials, participants were instructed to maintain central fixation and minimize eye movements and blinks; adherence was monitored by checking the EEG signal for stereotypical patterns of eye-movement-related artifact in frontal electrodes during the recording. Following each trial, a screen consisting of a fixation dot was shown for a randomly jittered inter-trial interval of 4-6 seconds. Participants were instructed to respond as quickly and accurately as possible, and completed a short practice block before the actual experiment.

Participants cared about their choices because they knew that a single randomly selected trial would be implemented for each recipient at the end of the experiment. If the participant had said yes on the selected trial, the recipient would have to eat at least three bites of the depicted food. If the participant said no, the recipient would not receive the food.

### EEG data acquisition and preprocessing

EEG data were collected using a 128-channel BioSemi ActiveTwo system (Biosemi B.V., Amsterdam, Netherlands) with active electrodes with sintered Ag-AgCl tips in fitted headcaps. Evoked brain potentials were digitized continuously at 512 Hz with default low-pass at 1/5 of the sampling rate. Two additional electrodes with a 4-mm sintered Ag-AgCl pallet were placed bilaterally on the mastoids for use as references in data import. Prior to the start of data collection, electrical offsets were verified to be between −20 and 20 μV across all channels.

Data preprocessing was performed offline using the EEGLAB toolbox (Delorme and Makeig, 2004) for Matlab (Mathworks, Natick, MA). Upon import into EEGLAB, data were re-sampled to 500 Hz and re-referenced to an average reference. To address DC offsets, linear detrending was performed on the continuous data. High-pass filtering at 1 Hz and notch filtering at 60 Hz was applied to remove slow voltage drifts and electrical noise, respectively, via a two-way least-squares FIR filter. Epochs were then extracted for a 2400 ms window around the onset of the food stimulus (−200 ms pre- to 2200 ms post-stimulus onset) for initial artifact rejection.

Although participants made an equal number of decisions for themselves and the two partners, subsequent division of the data into conditions by stimulus value or stimulus attribute (e.g., taste or health rating) produced unequal numbers of trials per condition due to individual variations in choice. Because traditional artifact rejection methods remove whole epochs, reducing the number of trials available to be averaged (Picton et al., 2000), use of these techniques could lead to even greater asymmetries in the distribution of trials by condition, potentially skewing the resulting average waveforms. Therefore, in lieu of traditional trial-based artifact rejection, we employed independent components analysis (ICA), a method that has been demonstrated to provide superior detection and removal for a variety of artefactual signals in EEG (Delorme et al., 2007; Uriguen and Garcia-Zapirain, 2015) without affecting the overall number of trials. Specifically, second-order blind identification (Belouchrani et al., 1997; Tang et al., 2005) (SOBI) was applied to “unmix” the EEG data into a sum of temporally correlated and spatially fixed components, which were then classified as task-related or artefactual (Harris et al., 2011). Whereas artifacts such as eye blink, muscle activity, and electrode noise tend to have highly localized scalp distributions and spectral peaks outside of typical EEG frequencies (e.g., 60-Hz line noise) without consistent time-locking to stimuli or responses, task-related cognitive components usually show meaningful dipolar scalp topographies, spectral peaks at typical EEG frequencies (e.g., 10-Hz “alpha”) and clear stimulus- or response-locking across the majority of trials. Only task-related components were projected back onto the scalp, allowing us to obtain artifact-corrected brain signals (Jung et al., 2000) from which 1600-ms stimulus-locked epochs (−100 ms pre-, 1500 ms post-stimulus onset) and 700-ms response-locked epochs (−600 pre-, 100 ms post-response onset) were extracted.

### EEG: Regression analysis

In our first analysis, we sought to characterize the time course of value signal computations, as identified by sensors and time windows in which neural activity correlated with the stimulus value of the foods. For each channel, evoked data for each trial were integrated over 50-ms windows from 100 to 1250 ms after stimulus onset, producing a matrix of EEG responses over 128 × 23 time windows in each participant. The response-locked data was likewise binned over 30-ms windows from −600 to 90 ms before response onset, resulting in 128 × 23 sensor-time window combinations in the response-locked data. For each analysis, the EEG responses for each participant across all 128 × 23 time windows were then entered into a linear regression model as in Equation 1:

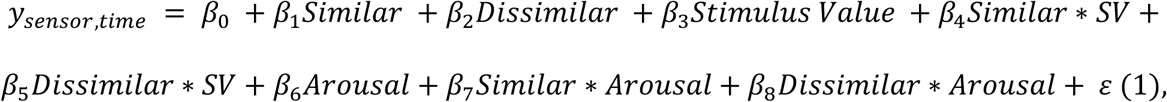

where *y_sensor, time_* consists of trial-by-trial data (in μV) for a particular sensor and time window, and β_0_ is the average activity in the sensor. The Similar and Dissimilar covariates have a value of 1 when the decision is for the specified recipient (*Similar* or *Dissimilar*, respectively) and 0 in all other cases. The Stimulus Value covariate encodes the value of the subject’s decision from 1 for Strong No to 4 for Strong Yes. The Arousal covariate, measuring the strength of preference regardless of valence (0 for Weak No or Weak Yes, 1 for Strong No or Strong Yes), was included to increase the statistical power of the linear model to identify activity associated with value (Litt et al., 2010). Finally, we computed separate covariates for the interaction of *Similar* and *Dissimilar* recipients with stimulus value and arousal, as indicated by Similar*SV, Dissimilar*SV, Similar*Arousal, and Dissimilar*Arousal. This linear regression analysis generated a set of estimated regression coefficients (i.e., beta map) for every sensor, time window, and subject. These maps were then aggregated into mixed-effect group estimates by computing one-sample t tests versus zero for each sensor and time window across all participants. The resulting *t*-statistic map allowed us to identify sensor and time window combinations in which neural activity was significantly correlated with stimulus value across participants. Results were corrected for multiple comparisons using an approximate permutation test in which the labels for all 600 trials were randomly shuffled 1000 times, and the resulting beta maps entered into one-sample t tests from which an empirical distribution of *t* values for each sensor and time window combination was then computed.

Next, we looked at how the weighting of taste and health attributes varied by recipient. We defined a time window of interest from 500 to 650 ms post-stimulus onset based on the time course of significant activity associated with the Stimulus Value covariate, which was also generally consistent with the timing of stimulus value effects in previous studies (Harris et al., 2011; Harris et al., 2013; Harris and Lim, 2016). Data in this time window was then inspected for the fully independent effects of recipient and attribute using a linear regression following Equation 2:

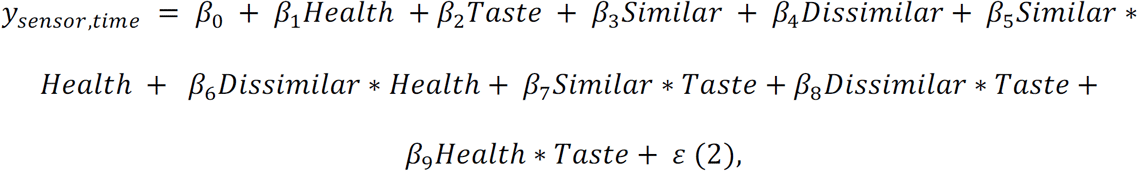

where the Health and Taste covariates correspond to the taste and health attribute ratings for each food item given by the individual participant, Health*Taste measures the interaction of the attribute ratings, and Similar*Health, Dissimilar*Health, Similar*Taste and Dissimilar*Taste reflect the interaction of a specific attribute with the identity of the recipient. Beta maps resulting from the linear regression in Equation 2 above were then compared using paired *t* tests of Similar*Taste – Similar*Health and Dissimilar*Taste – Dissimilar*Health to identify significant interactions of attribute by recipient. Specifically, sensors of interest (SOIs) were defined from the conjunction of significant sensors for the Similar*Taste – Similar*Health and Dissimilar*Taste – Dissimilar*Health comparisons within the predefined time window of interest (500-650 ms post-stimulus onset), and scalp topography and grand average waveforms were obtained for these SOIs.

### EEG: Bayesian source reconstruction

To localize activity associated with the Stimulus Value covariate to specific brain regions, we applied distributed source reconstruction using SPM8 (Wellcome Department of Imaging Neuroscience, Institute of Neurology, London, UK). In this approach, an empirical Bayesian algorithm (Friston et al., 2008) is used to model the cortical sheet as a series of hundreds of small dipolar patches, with the constraint that a common set of underlying sources account for the evoked responses across all participants (Litvak and Friston, 2008). In contrast to traditional dipole fitting, this method does not require *a priori* assumptions about the number or spatial locations of sources. Instead, source reconstructions were first computed across the entire trial to obtain models of activity in all potential sources and time points, followed by a more specific reconstruction of the time window of interest (WOI; 500 to 650 ms post-stimulus onset) defined on the basis of previous findings and the output of our mixed-effects ERP regression analysis.

Data from all participants were entered into the same source space using a “canonical mesh” based on the SPM template head model derived from the MNI brain. Sensors were coregistered with the MRI coordinate system using a generic template of the BioSemi 128-channel layout provided with the SPM software. Sensor locations were matched to the cortical mesh via an iterative alignment algorithm, and the source space was modeled using a boundary element model.

Our analysis employed a “localization of differences” approach (Henson et al., 2007), in which we searched for a set of neural sources representing different levels of the psychological variable of Stimulus Value. Difference waveforms were computed for each sensor and participant by multiplying the average waveforms for each condition by weights used to test for a linear (monotonically increasing) trend, and then summing across all weighted averages. To localize activity related to stimulus value computations, we used weights of (−3, −1, +1, +3) corresponding to the four levels of the Stimulus Value regressor, from Strong No to Strong Yes. To localize activity related to main effects of participant identity, contrast weights for Recipient corresponding to *Own, Similar*, and *Dissimilar* recipient choices were (−1, 0, 1) for the early 350 to 400 ms post-stimulus window and (−1, 1, 0) for the −300 to −180 ms pre-response window. Following reconstruction of the difference waveform for each participant, reconstructions for the WOI across participants were entered into a statistical analysis to identify statistically significant source estimates across individuals. In part due to the relatively high consistency of sensor locations across participants with the fixed cap layout used by this system, our initial analysis found large clusters of significant dipoles covering much of the brain, even at relatively rigorous family-wise-error (FWE) corrected statistical thresholds. Therefore, we chose a highly stringent threshold of *F* ≥ 100 to maximize separation of source estimates into distinct clusters while retaining significant dipoles. Results were visualized in terms of maximal intensity projection, focusing on cortical generators most likely to contribute to potentials recorded at the scalp (Cohen et al., 2011).

### Drift Diffusion Model Fitting: Time Varying Models

Our first modeling goal was to parameterize popular dual-process models of choice for others (i.e. anchoring and adjustment models) in a computational framework, in order to identify the most parsimonious set of parameters that accounted for patterns of choice and RT. Consistent with a growing body of work on value-based choices (Hutcherson et al., 2015; Tusche and Hutcherson, 2018) we assumed that choices can be captured using a multi-attribute variant of a drift diffusion model (DDM) (Smith and Ratcliff, 2004; Ratcliff and McKoon, 2008), in which noisy value signals accumulate over time and a choice is made when the accumulated signal crosses a pre-defined threshold for choice. In the simplest of these models, we assumed that this value signal (also known as the drift rate v) at each instantaneous time point *t* could be described as the linearly weighted sum of perceived tastiness and healthiness of the food on that trial, corrupted by Gaussian noise:

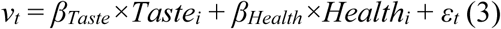

where *β_Taste_* and *β_Health_* represent the relative weights on Taste and Health, *Taste_i_* and *Health_i_* represent the values of Taste and Health attributes for the food shown on trial *i*, and *ε*_t_ is distributed ~N(0, .1). We assumed that the weights on Taste and Health could vary as a function of recipient (i.e., when deciding for the *Dissimilar* healthy partner, the weight on health might be higher than when deciding for oneself or for a *Similar*, unrestrained partner: *β_Health\Dissimilar_* > *β_Health\Similar_*).

The canonical version of the DDM assumes that *β_Taste_* and *β_Health_* remain constant throughout the course of a single decision. However, most dual-process models assume that participants make choices for similar others by anchoring first on their own preferences, and then serially adjusting away from this anchor using effortful perspective taking mechanisms. This suggests that *β_Taste_* and *β_Health_* might vary systematically over time, with early values resembling weights for one’s own self, and later values shifting to better reflect knowledge or assumptions about one’s partner. Although most descriptions of anchoring-and-adjustment models fail to specify precisely the temporal dynamics of this process, we attempted to capture their basic spirit using the following modifications to the standard DDM. First, we assume that the value signal at instantaneous time *t* can be described by the following equation

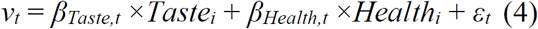

which is identical to Eq. 3, with the exception that *β_Taste_* and *β_Health_* are allowed to vary at each instantaneous time point *t*. Second, we assume that, when choosing for one’s *Own* self, *β_Taste_* and *β_Health_* are constant throughout the trial (i.e. *β_Taste|Own,0_* = *β_Taste\Own, t_* and *β_Health|Own,0_* = *β_Health\Own,t_*). Third, we assume that when choosing for others, participants begin with their own preferences (i.e. *β_Taste|Own,0_* = *β_Taste\Own, 0_* and *β_Health|Own,0_* = *β_Health\Own,0_*). However, these initial weights can begin to shift after a delay *T* to a new set of weights, at a rate *r* that determines how long the shift takes once it begins. We used these two parameters to capture the common notion in dual process models that perspective-taking might take time to implement, and that it should take longer the greater the difference in value between the old and new values. Thus, the full model when choosing for one’s *Own* self consisted of one fixed parameter (within-trial decision noise ε, which by convention is set to .1), and 4 parameters fitted to the data: weights on health and taste (*β_Taste\Own_* and (*β_Heaith\Owr_*), the choice-determining threshold (±*a*) and the non-decision time (*ndt*). Choices for others consisted of three fixed parameters (within-trial noise *ε* = .1, *β_Taste,0_* = *β_Taste\Own_* and *β_Health,0_* = *β_Health\Own_*) and six fitted parameters (new weights *β_Taste, t_*_> T_ and *β_Health,_*_t>T_, ±*a*, *ndt*, the transition delay *T* and the transition rate *r*).

Unfortunately, no known analytical solution to the DDM exists with time-varying drift rate parameters, rendering standard approaches to model-fitting and parameter estimation intractable. Thus, we employed a maximum-likelihood numerical simulation method similar to that employed in previous work on social decision-making (Hutcherson et al., 2015), with some important differences to speed up computation time (for details, see Tusche and Hutcherson, 2018). For each participant in each condition (*Own, Similar, Dissimilar*), we began by first simulating probability distributions of choices and response times for different combinations of parameters. We then used these probability distributions to identify the likelihood of the observed data for each parameter combination, and selected the parameter combination with the highest likelihood. Because this numerical simulation method takes a considerable amount of time that scales exponentially with the number of parameters and parameter values, we limited computational time by exploring a discrete set of values for each parameter. For the *Own* condition, we explored the following parameter values: threshold *a* = [.08, .1, .12, .14], non-decision time *ndt* = [.2, .3, …, .7], and *β_Taste_* = *β_Health_* = [−.02, −.01, −.005, 0, .005, .01, .02, ….14]. Parameter values for the Similar and Dissimilar partner conditions were identical except that the fitted values of *β_Taste_* and *β_Health_* represented the post-adjustment weights on Taste and Health after shifting away from the egocentric *Own* weights (estimated from the *Own* condition), and we included two additional time parameters to describe this shift: a delay parameter *T* = [0, .1, ., .5] sec, and a rate parameter *r* = [1.2, 2.4, 4.8, 9.6, 1000] describing how quickly old values shifted to new values following the delay. For example, a rate *r* = 1.2 suggests that it takes 1 second to shift 1.2 value units, or, equivalently, 250ms to shift .4 units. A rate *r* = 1000 indicates an instantaneous shift.

Using these values, we performed model fitting and model selection in the following manner. We first estimated the best-fitting parameter values for each participant’s choices in the *Similar* and *Dissimilar* partner conditions separately, estimating the full model with six free parameters. We then estimated a reduced, four-parameter model with only base parameters allowed to vary, *T* fixed at 0 (i.e. no early egocentric bias) and *r* fixed at 1000 (no gradual shift). Then, for each subject, we calculated the Bayesian Information Criterion (BIC) for each model based on the estimated likelihood of the data using the best-fitting parameter values. The winning model was determined based on which model had the lowest BIC value for the largest number of subjects, as well as the lowest BIC value when summing over all subjects. As we describe in greater detail in the results section below, this procedure identified the four-parameter model as the most parsimonious model to account for the data.

### Drift Diffusion Model Fitting: Constant Value Models

Because the analyses described above found no evidence for a time-varying adjustment process, all subsequent DDM estimation used canonical versions of the DDM, and was performed using a recently developed software package freely available in Python (Wiecki et al., 2013). Models were estimated using a Bayesian hierarchical framework, with Markov chain Monte Carlo (MCMC) sampling methods employed to estimate a joint posterior distribution of the model parameters. Hierarchical Bayesian estimation allows individual participant estimates to be constrained by a group distribution, but also to vary to the extent that their data demonstrate separation from the data of others. Estimation used non-informative priors: all priors were uniform distributions over large intervals of possible parameter values.

Several different DDM specifications were fit to the data. All of the DDM specifications assume the drift rate is a linear sum of weighted Taste and Health ratings, where the weights are allowed to vary by condition (*Own*, *Similar*, *Dissimilar*):

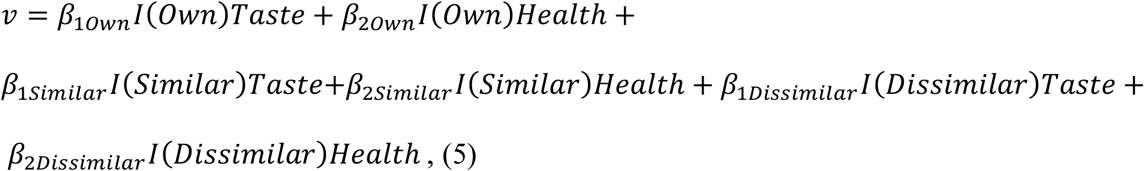

where I(Recipient) is an indicator function for each of the three conditions. The first model (M1) only allowed drift rate, as specified in Equation 3, to vary by task condition. All other parameters (*ndt,a,z*) were allowed to vary by subject, but did not vary within subject. The second model (M2) included drift rate as in M1, but also allowed the barrier parameter to vary with task condition, according to

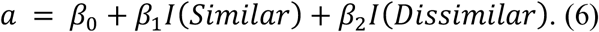

This model allowed for the possibility that, in addition to weighting attributes differently by recipient, individuals may also employ different thresholds depending upon the recipient. A third model (M3) considered the same drift rate as in M1, but also allowed the starting point, or bias, parameter *z* to vary with task condition, similar to Equation 4. This model did not allow barrier to vary with task condition. A fourth model (M4) allowed drift rate, barrier, and starting point to vary within participants. In all DDM specifications, the non-decision time parameter *ndt* was modeled to vary across participants, but not within participants.

We used two methods to compare fits of the various DDM specifications. First, we used the Deviance Information Criterion (DIC), a flexible measure for goodness-of-fit in hierarchical Bayesian models (Spiegelhalter et al., 2002). The DIC combines a measure of deviance (i.e., lack of fit) with a penalty for model complexity. A lower DIC represents a better model fit, and differences in DIC greater than 10 are typically thought to reflect significant differences (Spiegelhalter et al., 2002). Second, we compared observed data to data generated using random samples from the sampled posteriors (Wiecki et al., 2013). For each of the four models, 100 samples for each experimental subject were created. The squared error between each simulation and observed data was computed for both choices and RT, with the mean squared error (MSE) computed across the 100 runs. These statistics are reported for each model in Table 1.

**Table 1:** DDM Specifications Summary

| Parameter<br>w/ Trialtype | Model# | DIC | MSE Choice | MSE RT Yes | MSE RT No |
| --- | --- | --- | --- | --- | --- |
| v | M1 | 46384.2 | 0.0137 | 0.0537 | 0.0694 |
| v,a | M2 | 46318.6 | 0.0135 | 0.0532 | 0.0689 |
| v,z | M3 | 46522.1 | 0.0137 | 0.0559 | 0.0655 |
| v,a,z | M4 | 46415.6 | 0.0137 | 0.0538 | 0.0682 |

**Note:** v = drift rate, a = threshold, z = starting point.

Model convergence was assessed using the Gelman-Rubin (G-R) 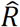 statistic (Gelman and Rubin, 1992). The 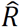 statistic compares within-chain and between-chain variance of different runs of the same model. Perfect convergence would be demonstrated with an 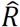 equal to 1. The 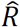 statistic was computed for all of the model parameters based on five runs of the model.

Another advantage of Bayesian estimation is that it is straightforward to compare differences in parameter estimates. Significance inference is possible using the posterior distributions generated from sampling. To compare coefficients to the null hypothesis (β = 0) the percentage of the N=10000 samples greater than zero can be assessed. Similarly, to determine whether or not a regression coefficient is greater than another, the percentage of the difference between the N=10000 samples that is greater than zero can computed.

### Drift Diffusion Model Fitting: Models Using ERP Data

In addition to the models M1 through M4, we also estimated a DDM using individual trial ERP components, which was identical to M2 in all ways except for additional regressors on the drift rate based on the ERP data (Equation 5). Specifically, as described above under “EEG: Regression Analysis”, we defined SOIs from the conjunction of significant sensors for Similar*Taste – Similar*Health and Dissimilar*Taste – Dissimilar*Health in the predefined time window from 500 to 650 ms post-stimulus onset. Within this SOI set and time window, we extracted average ERP amplitudes (in μV) separately for each participant and trial. Trial-by-trial ERP amplitudes were entered into a model with seven additional regressors beyond M2: one to measure the basic effect of the average trial-by-trial stimulus-locked ERP data on drift rate, and six to determine how trial-by-trial ERP data fluctuated with each of the decision targets and attributes (*Own, Similar, Dissimilar* for Health and Taste).

A similar procedure was followed to estimate the effect of response-locked ERP on the decision threshold. Trial-by-trial ERP amplitudes (in μV) were averaged across sensors showing a significant effect of *Similar* recipient identity between −300 and −180 ms pre-response. Along with behavioral choice and RT data, ERP amplitudes were entered into a model identical to M2 except for an additional regressor on the decision threshold to measure the effect of the trial-to-trial fluctuations in the ERP data (Equation 6). All estimation and statistical inference followed the same procedures outlined in the previous section.

### Post-Hoc Analysis: Individual Differences in Attribute Re-weighting by Neural Encoding of Identity

As we describe in greater detail below, we found neural responses that distinguished choices for the *Dissimilar* recipient relative to one’s *Own* self from approximately 350-400ms, localized to regions implicated in social cognition including the superior temporal sulcus (STS). At the same time, we observed that weights given to the Health and Taste attributes during choice varied as a function of recipient identity, with the largest differences in weights observed when comparing the *Dissimilar* other to the *Own* self. Taken together, these results suggest that neural responses from 350-400 ms may reflect a perspective-taking process that influences attribute re-weighting when deciding for others. If so, we speculated that the strength of this early ERP response should predict how much an individual shifts their weights on taste and/or health when choosing for others (particularly the *Dissimilar* other) compared to their *Own* choices. To test this idea, we correlated each individual’s neural response to recipient identity with the magnitude of their shifts in attribute weighting for each recipient.

To compute individual differences in the ERP response, we extracted average amplitudes between 350-400 m post-stimulus onset within the set of sensors showing a significant effect of *Dissimilar* recipient identity for this time window. Then, for each individual participant we calculated the difference in average ERP amplitudes separately for *Similar* vs*. Own* and *Dissimilar* vs*. Own* conditions. To compute individual differences in attribute weighting, we used the drift rate weights on Health and Taste attributes estimated from Model 2, Equation 5, to compute the following contrasts: 1) TASTE*_Similar vs. Own_*: change in Taste weight, *Similar* vs. *Own* trials (i.e., β_1Similar_ − β_1Own_); 2) TASTE*_Dissimilar vs. Own_*: change in Taste weight, *Dissimilar* vs. *Own* trials (i.e., β_1Dissimilar_ − β_1Own_); 3) HEALTH*_Similar vs. Own_*: change in Health weight, *Similar* vs. *Own* trials (i.e., β_2Similar_ − β_2Own_); 4) HEALTH*_Dissimilar vs. Own_*: change in Health weight, *Dissimilar* vs. *Own* trials (i.e., β_2Dissimilar_ − β_Own_). We then computed Spearman’s rank-order correlations between ERP*_Similar vs. Own_* and TASTE*_Similar vs. Own_*; ERP*_Similar vs. Own_* and HEALTH*_Similar vs. Own_*; ERP*_Dissimilar vs. Own_* and TASTE*_Dissimilar vs. Own_*; and ERP*_Dissimilar vs. Own_* and HEALTH*_Dissimilar vs. Own_*.

## Results

In this experiment, we investigated the neural and computational bases of decisions for self and others by measuring behavioral and ERP responses as participants with unrestricted diets made food choices for themselves and two partners: a similar partner with unrestricted preferences, and a dissimilar self-identified healthy eater (*Own* self, *Similar* other, and *Dissimilar* other conditions, Figure 1). First, participants rated 250 different appetitive snack foods on their perceived tastiness and healthiness. Second, for a subset of 200 foods selected to vary in taste and health, they were asked to decide whether they would select the pictured food for consumption after the experiment, making these choices once for each of the three recipients. To ensure that choices were incentive compatible, at the end of the study, one random trial was selected for each recipient and the participant’s choice on these trials was realized.

To characterize the neural and behavioral correlates of this social decision scenario, we performed a series of analyses as detailed in the following sections. First, we analyzed the behavioral choice and RT data to verify that participants were taking recipients’ preferences into consideration when making their decisions, and to replicate patterns previously associated with egocentric correction mechanisms. Next, we examined the stimulus-locked ERP data to find neural signals associated with stimulus value computations, further assessing how taste and health attributes were incorporated for each recipient during value integration. To determine how neural activity differed as a function of recipient, we compared ERP responses in each condition, time-locked to either the stimulus or the keypress response.

Although these analyses gave us a set of ERP components that correlated with different aspects of the decision process, they were blind to the specific cognitive operations indexed by these neural markers. Thus, to shed further light on the nature of these signals, we developed a set of novel multi-attribute computational models of choices for self and others based on extensions of the drift diffusion model (Ratcliff et al., 2016) and applied it to our event-related data. The primary assumption of these models is that weights on attributes like taste and health shape the evidence accumulated in favor of accept or reject responses. These weights can vary, not only as a function of choice recipient, but also as a function of time (i.e., biased toward automatic, egocentric defaults early on and then shifting over time with the application of controlled perspective taking). To test this latter assumption, we first implemented computational models in which the construction of value was allowed to vary systematically over time within a decision, and used formal model comparison and selection procedures to test whether and when this shift might occur. Second, having identified the simplest behavioral computational model to account for patterns of choice and RT, we then attempted to link trial-by-trial fluctuations in neural response to trial-by-trial variation in specific cognitive parameters such as the rate of evidence accumulation and threshold for response. Finally, applying our neurocomputational model to the behavioral data, we ran a series of simulation exercises demonstrating that distributions of choice and RT which appear consistent with default egocentric biases and slow effortful correction models of decision making can in some cases be explained without a correction mechanism, and assuming only relatively egocentric attribute weighting and value integration.

### Behavior: Observed choice and RT

To verify that participants made inferences consistent with their partner’s preferences, we first examined whether recipient identity influenced the likelihood of making a healthy choice (i.e., accepting/rejecting items rated high/low on health, Figure 2A). As expected, participants made healthier choices when choosing for the *Dissimilar* “healthy eater” compared to choosing either for *Own* self (paired-t_37_ = 9.19, *P* = 5×10^−9^) or the *Similar* other (paired-t_37_ = 10.49, *P* = 1×10^−12^). Participants made marginally *fewer* healthy choices for the *Similar* other (paired-t_37_ = 1.87, *P* = .069).

**Figure 2.**
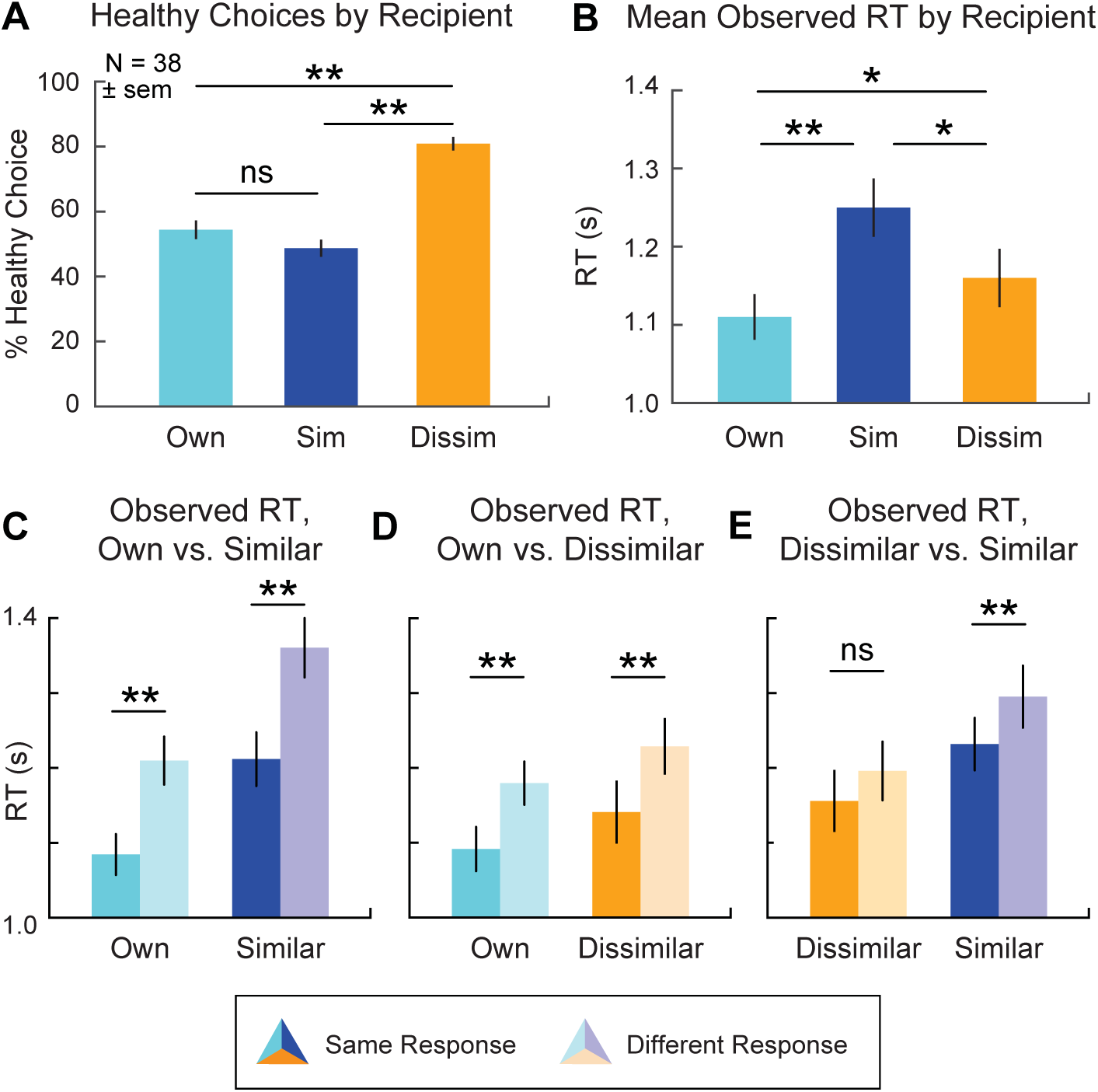
Behavioral results. (A) Percentage of healthy choices for each recipient. (B) Average observed RT by recipient. (C-E) Observed RT for trials separated on the basis of whether, given the same food option, the decision maker chose the same for (C) *Own* vs. *Similar*, (D) *Own* vs. *Dissimilar*, and (E) *Dissimilar* vs. *Similar*. Thus, *Own-Similar*:Same Response reflects RT for trials in which the choice was for *Own* self, and the response was the same as for the *Similar* other, whereas *Own-Similar*:Different Response reflects RTs on those trials where the response was different from the comparable choice for the *Similar* other; and likewise for the other comparisons.

We also asked how often participants made the same choice for themselves and their partner, one way of testing for egocentric responses. Participants made the same accept or reject choice for the *Similar* other as for themselves on 72±11% of trials, but chose the same significantly less often (60±17% of the time) when choosing for the *Dissimilar* other (paired-t_37_ = 3.69, *P* = .0007). These results suggested that participants tend to choose similarly for both recipients, with a more pronounced bias for the *Similar* other.

Next, we examined RTs for evidence that decisions for others might require additional computational resources. We analyzed behavior in two ways: average RT across all choices, and RT as a function of choice similarity between self and other. In line with the idea that choosing for others is more computationally costly, we observed significant differences in mean RTs depending on the recipient’s similarity (Figure 2B). Participants were fastest when choosing for themselves (mean = 1.11 s, SD = 0.18) and slowest when choosing for the similar other (paired-t_37_ = 7.59, *P* = 5×10^−9^), with choices for the dissimilar partner lying in between and significantly different from the other two conditions (both paired-t_37_ > 2.17, both *P* < .036).

A critical finding from previous research is that individuals take longer to choose for similar others when making choices that differ from their own preferences, but that this effect is reduced when choosing for dissimilar others (Epley et al., 2004; Tamir and Mitchell, 2013). As seen in Figure 2C-E, we replicated this pattern, observing longer RTs when choices diverge from *Own* preferences, for both *Similar* (different choice: mean = 1.37 s, SD = 0.25; same choice: mean = 1.22 s, SD = 0.28; paired-t_37_ = 7.90, *P* = 2×10^−9^) and *Dissimilar* (different choice: mean = 1.23 s, SD = 0.26; same choice: mean = 1.14 s, SD = 0.23; paired-t_37_ = 3.35, *P* = .002) conditions. This effect was marginally more pronounced in choices for the *Similar* partner compared to the Dissimilar partner (paired-t_37_ = 1.75, *P* = .089), in keeping with prior work (Tamir and Mitchell, 2013).

Intriguingly, however, we also found a slowing effect when making choices for one’s self that diverged from the choices made for partners (*Own* vs. *Similar:* mean RT_Different_ = 1.22±0.20 s, mean RTS_ame_ = 1.09±0.17 s, paired-t_37_ = 6.27, *P* = 2×10^−5^; *Own* vs. *Dissimilar:* mean RT_Different_ = 1.23±0.26 s, mean RT_Same_ = 1.14±0.23 s, paired-t_37_ = 3.35, *P* = .002). This analysis has not been reported previously, to our knowledge, and raises important questions about the most parsimonious explanation of RT differences for divergent choices, as we discuss in greater detail with respect to our computational simulation results below.

### ERP: stimulus value integration

Given that participants adjusted their choices to reflect the recipient’s preferences, a further question is when and how relevant value signals emerge in the brain. Although fMRI studies have identified overlapping regions of VMPFC representing value for both oneself and others (Nicolle et al., 2012; Janowski et al., 2013), some dual-process models suggest that these signals should take longer to construct for others. Therefore, the high temporal resolution of ERP is of crucial importance in determining the time course of stimulus value integration when choosing for oneself versus others. We began by identifying neural correlates of stimulus value computation, which previous neurophysiological studies of choice for oneself have identified between 400 and 650 ms after stimulus onset, localized to regions including the VMPFC (Harris et al., 2011; Harris et al., 2013; Harris and Lim, 2016).

We applied a linear regression analysis to the ERP data in order to identify neural activity correlated with the decision made in each trial (1 = Strong No to 4 = Strong Yes) across participants. As shown in Figure 3A, ERP data was binned over 50-ms windows from 100 ms to 1200 ms after stimulus onset, for all 128 sensors grouped by scalp location. Data were permutation-corrected to account for multiple statistical comparisons. This analysis revealed significant main effects of stimulus value, regardless of recipient, at two major time windows leading up to the time of choice. Generally consistent with the time range of stimulus value effects reported in previous studies, the earliest major cluster of value-related activity occurred from approximately 500 to 650 ms after stimulus onset (Figure 3A, green box), with additional activity in roughly the same set of sensors re-emerging from 700 to 850 ms post-stimulus onset (Figure 3A, blue box). Distributed source reconstruction of scalp topography for these two time windows localized the stimulus value signal to regions including VMPFC, with a particularly pronounced localization to VMPFC in the 500-650 ms time window (Figure 3B, circled). Although the coarse spatial resolution of ERP precludes detailed specification of sources, the sources found here closely resemble those from previous fMRI studies (Figure 3B, top inset), as well as prior ERP localizations (Figure 3B, bottom inset) within the same latency range (Harris et al., 2013; Harris and Lim, 2016).

**Figure 3.**
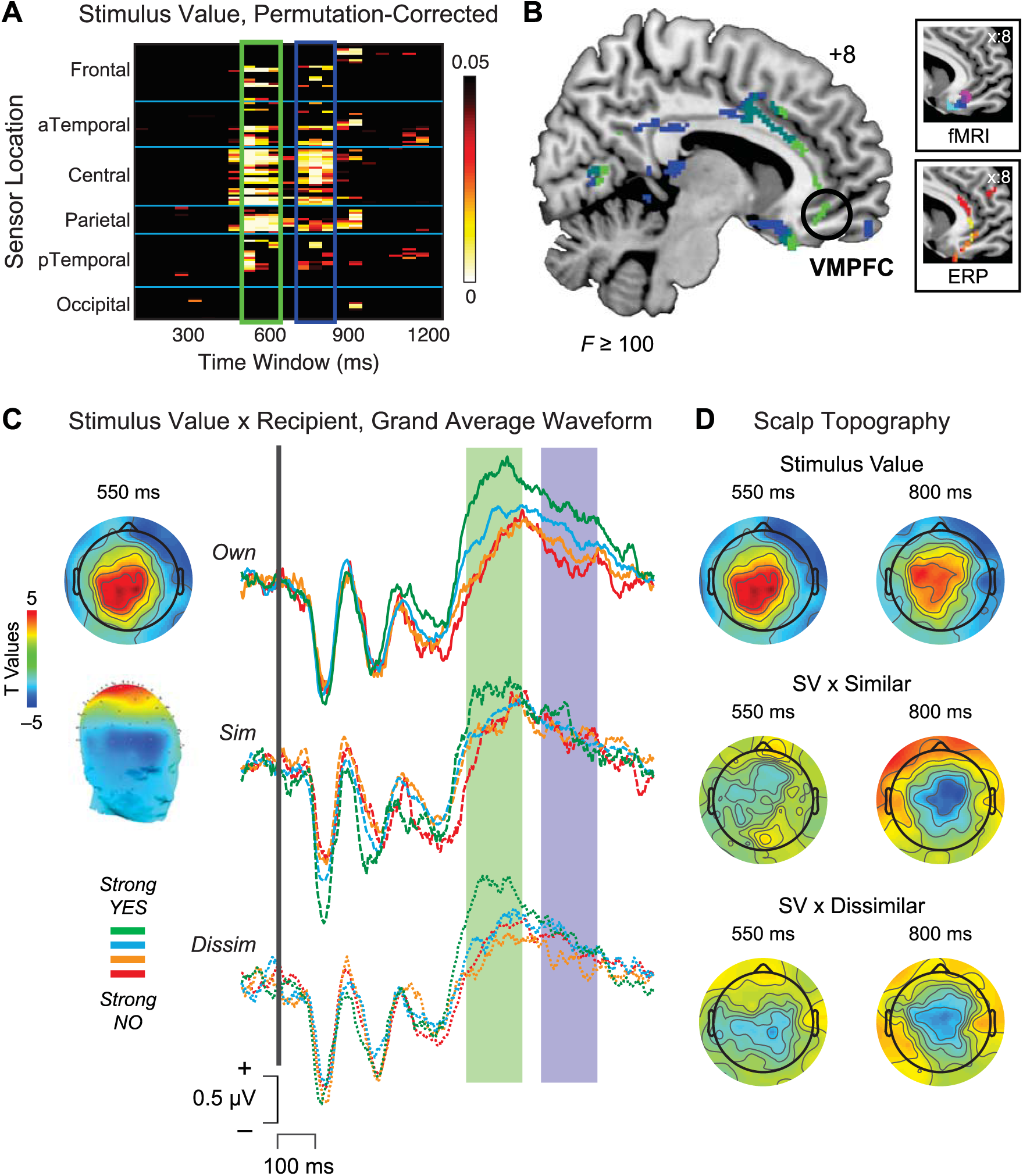
ERP analysis of stimulus value integration by recipient. (A) Heat map of significant *P* values associated with stimulus value, showing significant effects from 500 to 650 ms (green box) and 700 to 850 ms (blue box) after stimulus onset. (B) Source reconstructions from 500 to 650 ms (green) and 700 to 850 ms (blue) overlaid on a representative brain image. Particularly during the earlier 500-650 ms window, stimulus value activity was localized to regions including VMPFC (circled). Inset, top: Spherical masks based on peak coordinates from three neuroimaging studies [blue (Plassmann et al., 2007), magenta (Hare et al., 2009), cyan (Litt et al., 2010)]. Inset, bottom: Source localization of stimulus value from approximately 450 to 600 ms post-stimulus onset in two ERP studies [red: (Harris et al., 2013), yellow: (Harris and Lim, 2016)]. (C) Stimulus value integration by recipient. Topographic scalp distribution (left) and average waveforms for the linear ordering of stimulus value (red: Strong No; orange: Weak No; cyan: Weak Yes; green: Strong Yes) in the 500-650 ms window (shaded green box) and 700-850 ms window (shaded blue box), plotted separately for each recipient (*Own*: solid, *Similar*: dashed; *Dissimilar:* dotted). (D) Comparison of scalp topography at 550 ms (left) and 800 ms (right) post-stimulus for the main effect of stimulus value (top), the interaction of stimulus value with *Similar* recipient (middle) and the interaction of stimulus value with *Dissimilar* recipient (bottom) revealed significant reductions in stimulus value signals for the two other recipients during the late window, in line with average waveform data in (C).

Given the overlapping sensor distributions and source localizations for the earlier and later time periods, what differentiates neural value integration during these two windows? To address this question, we examined stimulus value computations as a function of recipient identity using the conjunction of significant sensors 500-650 ms and 700-850 ms after stimulus onset (Figure 3C). Grand average waveforms from central sensors (Figure 3C, left) revealed a clear linear ordering from Strong No to Strong Yes for all recipients between 500 and 650 ms post-stimulus (Figure 3C, shaded green region). In contrast, activity in the later 700-850 ms window demonstrated a strong “self-relevance” effect (Grueschow et al., 2015): the strongest Stimulus Value responses were observed for one’s *Own* self, perhaps reflecting sustained attention to one’s own choices, with diminished responses for *Similar* and *Dissimilar* partners (Figure 3C, shaded blue region).

To confirm these effects statistically, we examined covariates representing the interaction of stimulus value with each recipient that were computed as part of the larger regression analysis (Eq. 1). Visualizing these effects at the scalp separately for time windows 550 and 800 ms post-stimulus onset (Figure 3D) revealed that, despite highly significant activity at central sensors for stimulus value in the earlier window (Figure 3D, top left), the interactions of stimulus value with *Similar* (Figure 3D, middle left) and *Dissimilar* (Figure 3D, bottom left) were largely non-significant, with *t*-values surviving a threshold of *p* ≤ 0.05 in 0 and 1 sensors, respectively. In contrast, during the later window (Figure 3D, right), interactions of recipient and stimulus value were significantly negative for both partners (*Similar:* 18 sensors, average *p* = 0.01; *Dissimilar:* 12 sensors, average *p* = 0.03), suggesting that stimulus value computations for others are diminished rather than enhanced during this late time window. Thus, we did not find stronger delayed value representations during decisions for others, as would be expected if participants required additional time to construct others’ preferences. Instead, later value responses more likely reflect sustained attention and/or arousal to choices with a direct impact on one’s own outcomes.

### ERP: attribute weighting by recipient

In line with the hypothesized role of VMPFC in context-dependent value integration (Hare et al., 2009), ERP value signals from roughly 500 to 650 ms post-stimulus have previously been shown to reflect increased weighting of health attributes when participants are incentivized to exercise dietary self-control (Harris et al., 2013), overlapping with the time window of significant stimulus value effects in our current data. Thus, we predicted that neural activity in the time window of 500-650 ms post-stimulus onset would show differential weighting on taste and health depending on the recipient of the choice.

To test this, we computed a second regression identifying sensors in which neural activity significantly correlated with the interactions of *Similar* and *Dissimilar* recipient by Taste and *Similar* and *Dissimilar* recipient by Health (Equation 2). The beta maps resulting from this regression were then compared using paired t-tests to find sensors where the relative weighting of taste (Figure 4A, left) and health (Figure 4A, middle) varied by recipient during our time window of interest from 500 to 650 ms after stimulus onset. Difference plots of scalp topography in this time window identified a set of central sensors (Figure 4A, right) for the interaction of taste and health in both *Similar* and *Dissimilar*, overlapping those associated with stimulus value in our initial analysis. Sensors of interest (SOIs) were formally defined via the conjunction of sensors associated with significant weighting on Taste – Health in both *Similar* and *Dissimilar* recipient conditions.

**Figure 4.**
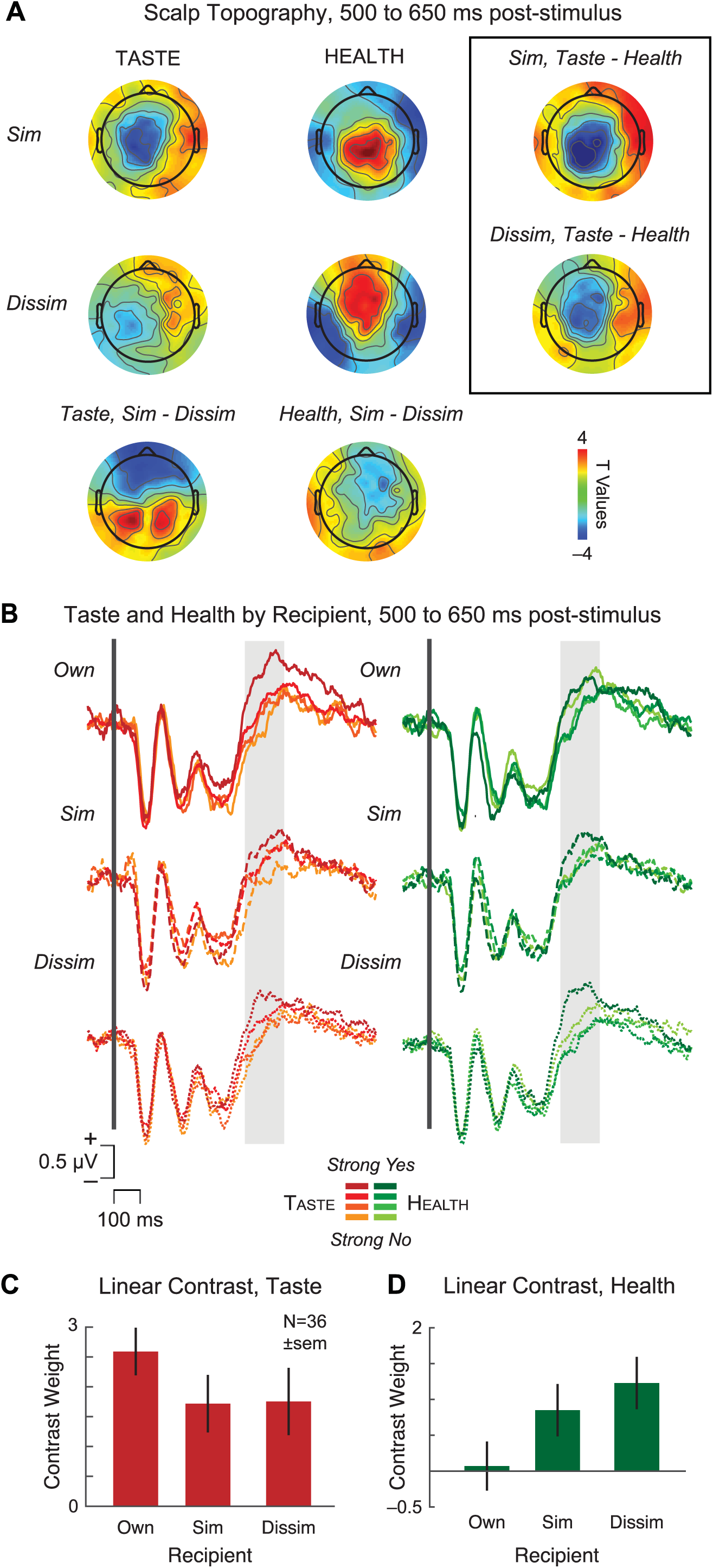
ERP analysis of attribute weighting by recipient. (A) Scalp topography during time window of interest from 500 to 650 ms post-stimulus revealed differences in neural weighting of taste ratings (left) and health ratings (middle) by recipient. Sensors of interest (SOI) were identified by taking the conjunction of sensors showing significant ERP activity for *Similar* and *Dissimilar* in the Taste – Health comparison (black box, right). (B) Average waveforms associated with the linear ordering of taste (left) and health (right) plotted separately for each recipient (*Own*: solid; *Similar*: dashed; *Dissimilar*: dotted). Orange: taste; Green: health. (C-D) Linear contrast weights for taste (C) and health (D) as a function of recipient.

To clarify the direction of the interaction defined by our conjunction analysis, we next examined grand average waveforms and linear contrasts extracted from the SOIs (Figure 4B-D) separately for taste and health attributes by recipient. In line with our predictions, the clearest parametric effect of taste was visible when deciding for one’s *Own* self versus the two partners (Figure 4B, left; Figure 4C), whereas the linear ordering of health showed the greatest effect for the *Dissimilar* healthy eater and negligible distinction by health for one’s *Own* self (Figure 4B, right; Figure 4D). We confirmed these observations statistically using a repeated-measures ANOVA on the linear contrast weights with Attribute (Taste/Health) and Recipient (Own/Similar/Dissimilar) as covariates, focusing on the interaction of Attribute x Recipient. We found a significant interaction of Attribute x Recipient (*F*(2,70) = 5.05, *p* = 0.009), reflecting a significant linear contrast (*F*(1,35) = 8.39, *p* = 0.006) driven by the opposing linear trends in neural weighting of taste and health attributes across recipients. These results support the idea that value integration from approximately 500 to 650 ms post-stimulus reflects differential attribute weighting for oneself and others.

### ERP: neural signals related to recipient identity

The findings described above bolster the idea that decisions for others recruit similar value computation mechanisms as decisions for oneself, on the same time scale, regardless of how dissimilar the recipient’s preferences may be. However, these results raise an additional question: how and when does the brain modify value computations to distinguish the intended recipient of the choice? To address this issue, we next looked for neural activity that differentiated recipient identity, using ERP data time-locked to both stimulus and response.

Some versions of a dual-process account suggest that inhibition of egocentric biases should emerge relatively late in the process and, because similar individuals more strongly activate egocentric biases that need correcting, such inhibition should be more pronounced for the *Similar* other (Tamir and Mitchell, 2013). In contrast, other research suggests attention and perspective taking mechanisms should be most pronounced for the *Dissimilar* other, whose preferences differ most markedly from one’s *Own* self (Silani et al., 2013). Given that we found a similar time course of value integration by recipient, we hypothesized that this perspective-taking and adjustment process might actually occur *early*, prior to or during the window of stimulus value computation (i.e., prior to 500 to 650 ms post-stimulus). Previous neuroimaging research has suggested that regions associated with social cognition, such as TPJ and superior temporal sulcus (STS), play a key role in directing value integration in VMPFC during social decisions (Hare et al., 2010; Hutcherson et al., 2015; Strombach et al., 2015). Therefore, we further predicted that any differences would localize to regions including TPJ.

To test these predictions, we computed a linear regression with separate dummy-coded indicator variables for *Similar* and *Dissimilar* recipient; the results of this analysis from 100 to 600 ms after stimulus onset, our *a priori* time window of recipient computation, are shown in Figure 5A. Although the covariate for *Similar* recipient (Figure 5A, left) was largely non-significant during this time window, we found significant ERP activity for the *Dissimilar* recipient from approximately 350 to 400 ms post-stimulus onset (Figure 5A, right). Average waveforms for these sensors and time window (Figure 5B) revealed a positive response for the *Dissimilar* condition relative to one’s *Own* self, localized to sources including posterior superior temporal sulcus (STS) and TPJ (Figure 5B, right). Consistent with previous studies (Saxe, 2006; Andrews-Hanna et al., 2010; Hare et al., 2010; Hutcherson et al., 2015), these results suggest a mechanism by which brain networks involved in mentalizing about others may communicate with valuation regions in order to adjust attribute weights during value integration.

**Figure 5.**
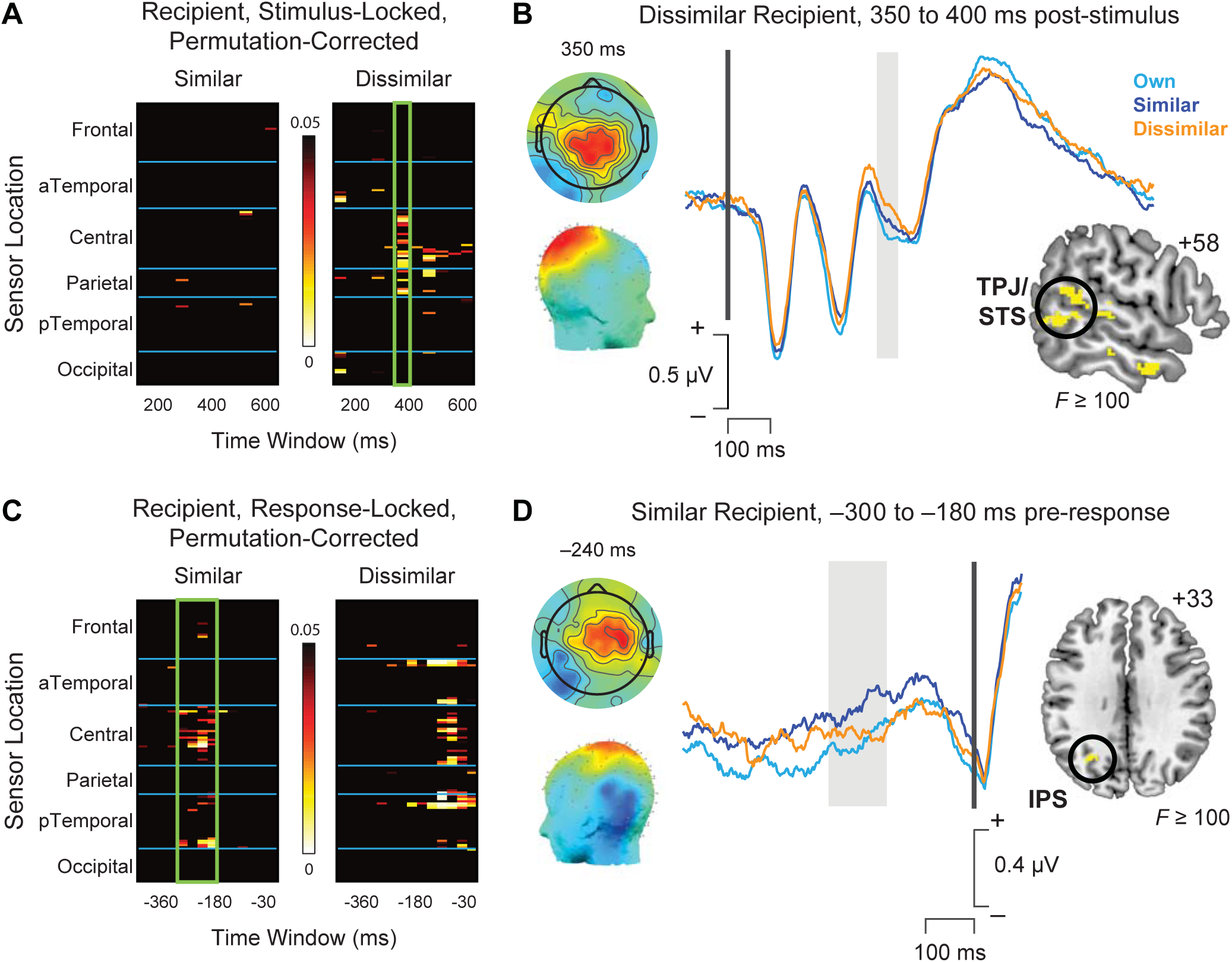
Differences in average ERP waveforms by recipient identity. (A) Heat map of significant *P* values showing stimulus-locked regression results for the main effects of *Similar* (left) and *Dissimilar* (right). Significant effects were visible for the *Dissimilar* recipient, but not the *Similar* recipient, at time windows including 350 to 400 ms after stimulus onset (green box). (B) Scalp topography (left) and average waveforms (right) associated with choice recipient identity (*Own* self: cyan, *Similar* other: blue, *Dissimilar* other: orange) 350-400 ms post-stimulus onset. Inset: Source reconstruction from 350-400 ms post-stimulus overlaid on a representative brain image localized this response to regions including the temporoparietal junction (TPJ) and superior temporal sulcus (STS). (C). Heat map of significant *P* values showing response-locked regression results for the main effects of *Similar* (left) and *Dissimilar* (right). Significant effects were visible for the *Similar* recipient, but not *Dissimilar* recipient, approximately 300ms to 180 ms prior to the response (green box). Later effects in the Dissimilar condition coincide with motor activity related to initiation of the key press response. (D) Scalp topography (left) and average waveforms (right) associated with choice recipient identity (*Own* self: cyan, *Similar* other: blue, *Dissimilar* other: orange). Inset: Source reconstruction from 300 to 180 ms prior to the response localized this differential response to regions including the intraparietal sulcus (IPS).

Although pre-value signals differentiated the *Dissimilar* other from both *Own* self and *Similar* other trials, we also sought to test predictions of several dual-process models that there may also be a late adjustment process, unique to choices for *Similar* others, that delays choices and corrects egocentric values to account for differing preferences. Although we did not find evidence for late value integration signals when deciding for *Similar* others (see “Stimulus value integration” results above), we nevertheless asked whether there was a main effect of recipient identity in late-emerging response-locked ERP signals, as might be predicted by some dual-process models. Intriguingly, a linear regression on ERP data time-locked to the keypress response revealed a significant main effect of *Similar* recipient from approximately −300 ms to −180 ms pre-response (Figure 5C), reflecting a more positive deflection for *Similar* relative to *Own* and *Dissimilar* (Figure 5D). To shed light on the nature of this process, we performed source localization from −300 to −180 ms pre-response. In this time window, the main effect of *Similar* recipient was predominantly localized to the intraparietal sulcus (IPS), close to an area previously identified as being more activated when choosing for others (Janowski et al., 2013). In contrast, significant main effects of *Dissimilar* recipient in this time window were largely confined to the period immediately prior to response (−120 to 0 ms pre-response), and thus are more likely to reflect the initiation of movement potentials associated with the motor response.

### Computational Modeling

As described above, our analysis of the time course of value integration suggests that a *single* valuation mechanism, localized to the VMPFC, computes stimulus values using weighted attribute integration for both self and others, with similar temporal dynamics. Yet the presence of both early- and late-emerging components in TPJ and IPS that were specific to choosing for others is fully consistent with dual-process models of choice in which perspective-taking mechanisms alter value computations to reduce egocentric biases. This raises an important question: what is the most parsimonious computational framework to account for both the behavioral and neural data?

To address this issue, we used the power of formal model comparison and selection procedures to identify the most parsimonious computational model of decision making for oneself versus others, and then explored how changes in parameter values across choices for self and others shed light on neural and behavioral responses. We began by specifying a multi-attribute variant of the DDM that allowed us to capture basic assumptions of dual-process accounts, which suggest that the construction of value should vary in systematic ways over the course of a decision. More specifically, we assumed that weights on taste and health attributes influence the overall drift rate (*v*), and that these weights should be egocentrically biased early on during a decision but shift gradually after a delay to a new set of weights that reflect a participant’s assumptions about the preferences of their partner after deliberation. We compared this model to a nested model in which, when choosing for others, participants simply begin at the outset with a set of weights that reflect their assumptions about the other person’s preferences, with no delay or gradual adjustment process (see Methods for details). Remarkably, for the *Similar* condition, the nested model with no time-varying parameters had a lower BIC value combining over all subjects (1.2159 × 10^5^) compared to the time-varying model (1.2189 × 10^5^), as well as a lower BIC value for 36 out of 38 individual subjects. Conclusions were largely similar using the more liberal Akaike Information Criterion (AIC) instead of BIC. We arrived at identical conclusions when examining choices for the *Dissimilar* other (total BIC of the simple model = 1.1487 × 10^5^, total BIC of the time-varying model = 1.1518 × 10^5^).

Having shown that a simple DDM provided a more parsimonious fit to the data, we then sought to determine how the values of different model parameters might change as a function of choices for *Own Self, Similar*, and *Dissimilar* others. Specifically, we implemented a multi-attribute version of the DDM with five parameters: weights on the taste and health attribute in the drift rate (*v*), threshold for choice (*a*), starting point bias (*z*), and non-decision time. We fit four different versions of the DDM to the data, in which drift rate, threshold, and starting point bias were allowed to vary depending on the choice recipient (Table 1). The best-fitting model, as assessed by both lower DIC values and lower MSE (M2; Table 1) suggested that, as expected, weights on health and taste varied significantly by recipient. On average, participants weighted both the health and taste attributes more similarly to themselves for the Similar individual than for the Dissimilar individual (Table 2). However, in all cases there were significant pairwise differences between attribute weights for different recipients. Taste ratings were weighted highest for *Own* self, somewhat less for the *Similar* other, and not at all for the *Dissimilar* other (all comparisons: 100% of the difference in posteriors > 0, i.e., all *P*-values = .000). Health ratings were weighted highest for *Dissimilar*, significantly less for the *Similar* other, and least for *Own* self (all comparisons significantly different, all *P* = .000).

**Table 2:** DDM posterior estimates and convergence for Model M2

| Parameter | Mean | Stdev |
| --- | --- | --- |
| Drift (v), Intercept | -1.982 | 0.055 |
| Drift (v), HealthOwn | 0.062 | 0.011 |
| Drift (v), HealthSimilar | 0.183 | 0.010 |
| Drift (v), HealthDissimilar | 0.660 | 0.011 |
| Drift (v), TasteOwn | 0.722 | 0.011 |
| Drift (v), TasteSimilar | 0.665 | 0.010 |
| Drift (v), TasteDissimilar | 0.007 | 0.011 |
| Barrier (a), Intercept | 2.006 | 0.047 |
| Barrier (a), Similar | 0.125 | 0.015 |
| Barrier (a), Dissimilar | 0.049 | 0.015 |
| Starting point (z) | 0.493 | 0.006 |
| NonDecisionTime | 0.357 | 0.026 |
| **Convergence** | **Mean** | **Stdev** |
| $\hat{R}$ | 1.0002 | 0.0002 |

**Note:** Statistics for parameter estimates reflect the mean and standard deviation of the N=10000 posterior samples of group-level parameters.

Although weights on taste and health attributes differed significantly on average during choices for both *Similar* and *Dissimilar* others compared to *Own* self, most models of choice assume that participants also show egocentric biases, using their own preferences when making choices for others (e.g., Devaine and Daunizeau, 2017; Tarantola et al., 2017). To test this possibility, we theorized that if people do use their own preferences to some degree when choosing for others, then individual differences in the weights estimated for Taste and Health attributes in *Own* choices should predict the weights estimated for these attributes when choosing for others, particularly when those preferences are relevant to the choice at hand. This prediction was robustly confirmed. Individual weights on taste for one’s *Own* self correlated strongly with weights on taste for the *Similar* other (*r*_36_ = .58, *P* = .0002) but not for the *Dissimilar* other (*r*_36_ = .07, *P* = .66). Individual weights on health for one’s *Own* self correlated with weights for both the *Similar* other (*r*_36_ = .57, *P* = .0002) and *Dissimilar* other (*r*_36_ = .48, *P* = .003). Thus, our best-fitting model strongly supports the existence of an egocentric bias when making choices for others.

Intriguingly, the best-fitting model also suggested significant variation in the threshold parameter *a* as a function of choice recipient. Participants adopted greater caution when choosing for others compared to themselves, with the highest threshold associated with choosing in the *Similar* other condition. We speculate that these thresholds are adjusted to accommodate greater uncertainty about the appropriate choice, and that this uncertainty may have been highest for the *Similar* other, a point we return to in the Discussion. In contrast to attribute weights and threshold, starting point (i.e. response bias) did not significantly differ from an unbiased value of 0.5 (11.9% of posterior > 0.50, i.e., *P* = .119) and was similar for all recipients.

To confirm the ability of the DDM to successfully capture both observed choice and RT data, posterior predictive simulations (Wiecki et al., 2013) were generated using 100 randomly selected samples from the posterior, effectively generating 100 synthetic datasets. Posterior predictive checks using this simulated data confirmed the ability of the model to recreate acceptance rates and RTs at both the group and individual level. Pearson correlations between the true data and synthetic data for participant-level average data were for high for both acceptance rates (*Own*: 0.863, *Similar:* 0.750, *Dissimilar:* 0.755) and RT (*Own*: 0.934, *Similar:* 0.921, *Dissimilar:* 0.942). Importantly, convergence statistics suggested stable parameter estimates across all parameters in the hierarchical model (Table 2).

### Combined ERP-model results

We next sought to validate our model not only through behavior, but by linking specific ERP components to specific model parameters. We predicted that if value signals in the VMPFC measured on-line during decision making between 500-650ms represent the most proximal input into the evidence accumulation process, fluctuations in this signal should be directly related to variation in the model-estimated influence of taste and health on the drift rate, *even after accounting for behavioral ratings* collected after the fact. To test this prediction, we entered trial-by-trial stimulus-locked ERP data from our SOIs and 500-650 ms time window of interest (Figure 6A) into the M2 specification for the DDM, interacted with both Health and Taste ratings. Note that stimulus-locked signals in the VMPFC were identified based on their association with health and taste ratings collected independently of the choice, and are here assumed to be an *input* into the accumulation process, and not an output of it.

**Figure 6.**
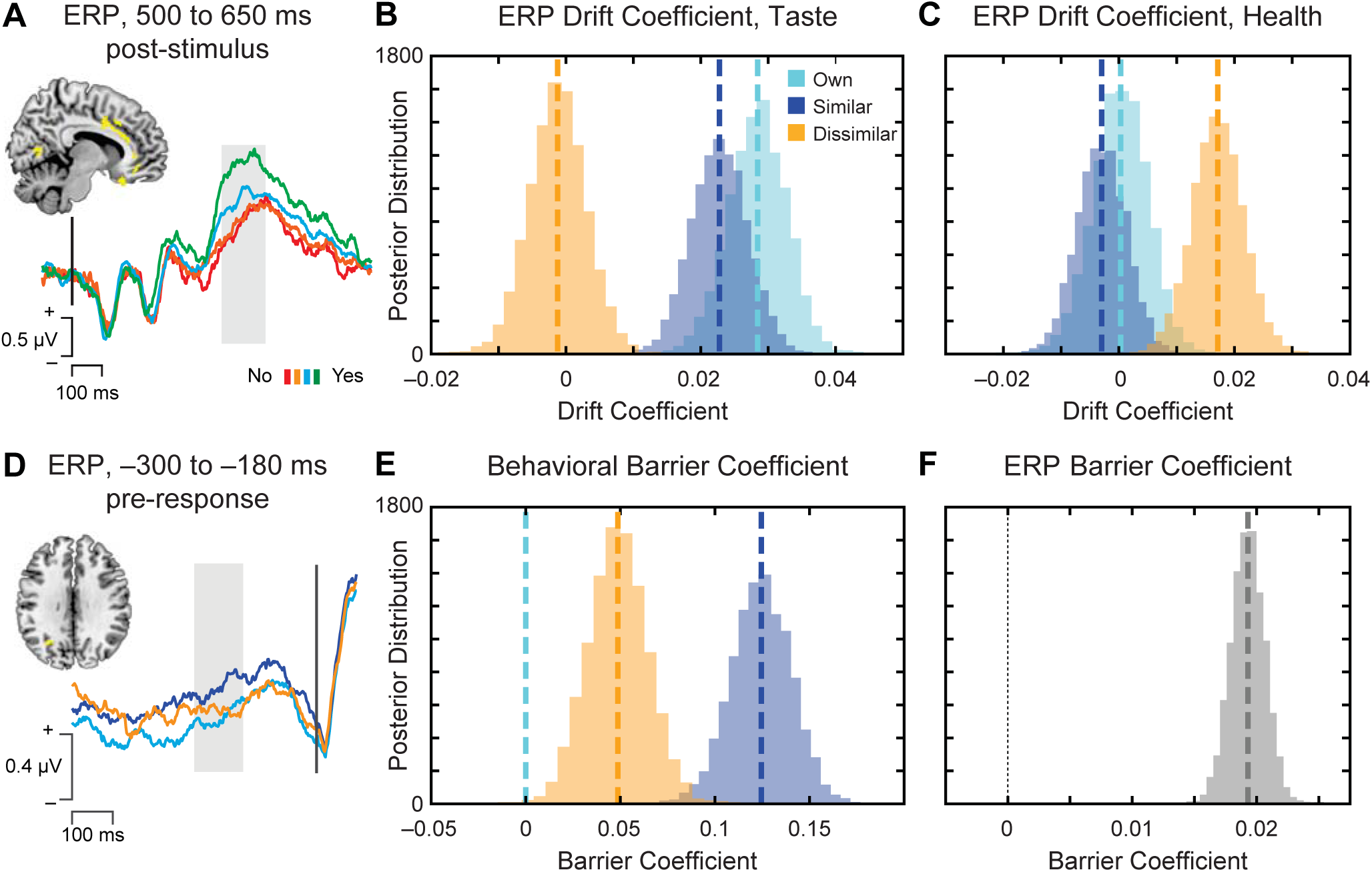
Hierarchical Bayesian parameter estimates of the influence of neural response on model parameters. (A-C) Trial-by-trial fluctuations in the ERP in sensors-of-interest (SOIs) for the value-related ERP from 500-650ms (A) were linked with significantly greater influence on the drift rate *v* of the taste attribute when choosing for *Own* self and *Similar* other (B) and the health attribute when choosing for the *Dissimilar* other (C), suggesting that this response relates to evidence accumulation in the computational model. (D-F) Trial-by-trial fluctuations in the response-locked ERP component (D) differentiated choices for the *Similar* other to a greater extent than *Dissimilar* or *Own* choices, consistent with computational model-fitting of the behavioral barrier parameter (E), and likewise demonstrated a significant influence on the barrier for response (F).

As predicted, during choices for both *Own* self and *Similar* other, the amplitude of the ERP value response predicted a significantly higher influence of Taste on the drift rate when choosing for *Own* self and *Similar* other (Figure 6B, 100% of the posteriors for the interaction > 0, i.e., *P* = .000), but not when choosing for the *Dissimilar* other (P = .380). *Own* and *Similar* Taste*ERP effects were not significantly different from each other (*Own* > *Similar*, P=. 118), but were both significantly greater than *Dissimilar* (*P* = .000). By contrast, the amplitude of the ERP response significantly enhanced the influence of the Health attribute when choosing for the *Dissimilar* other (P = .000), and this effect was significantly greater than for *Own* (P=.001) or *Similar* (*P* = .000) choices (Figure 6C). Notably, the addition of parameters reflecting ERP activity to the drift rate did not substantially alter the parameter estimates derived from behavioral data (Table 3), suggesting that the neural data provide an additional, unique source of information to our computational model. These results strongly support the idea that VMPFC value signals from 500-650ms contribute to choice by influencing the accumulation of attribute evidence toward a choice.

**Table 3:**
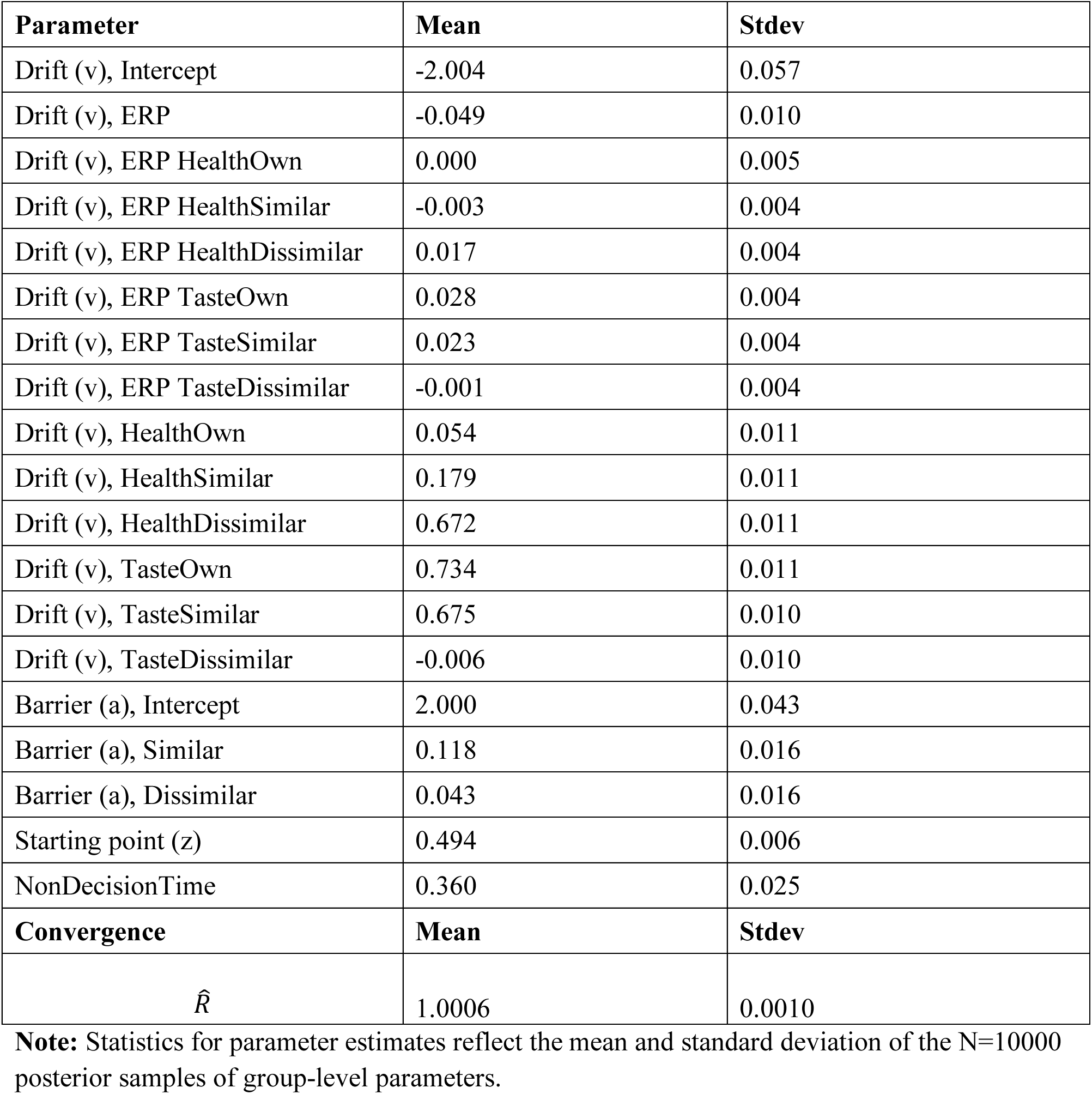
DDM posterior estimates and convergence for Model M2 with stimulus-locked ERP data.

| Parameter | Mean | Stdev |
| --- | --- | --- |
| Drift (v), Intercept | -2.004 | 0.057 |
| Drift (v), ERP | -0.049 | 0.010 |
| Drift (v), ERP HealthOwn | 0.000 | 0.005 |
| Drift (v), ERP HealthSimilar | -0.003 | 0.004 |
| Drift (v), ERP HealthDissimilar | 0.017 | 0.004 |
| Drift (v), ERP TasteOwn | 0.028 | 0.004 |
| Drift (v), ERP TasteSimilar | 0.023 | 0.004 |
| Drift (v), ERP TasteDissimilar | -0.001 | 0.004 |
| Drift (v), HealthOwn | 0.054 | 0.011 |
| Drift (v), HealthSimilar | 0.179 | 0.011 |
| Drift (v), HealthDissimilar | 0.672 | 0.011 |
| Drift (v), TasteOwn | 0.734 | 0.011 |
| Drift (v), TasteSimilar | 0.675 | 0.010 |
| Drift (v), TasteDissimilar | -0.006 | 0.010 |
| Barrier (a), Intercept | 2.000 | 0.043 |
| Barrier (a), Similar | 0.118 | 0.016 |
| Barrier (a), Dissimilar | 0.043 | 0.016 |
| Starting point (z) | 0.494 | 0.006 |
| NonDecisionTime | 0.360 | 0.025 |
| **Convergence** | **Mean** | **Stdev** |
| $\hat{R}$ | 1.0006 | 0.0010 |

**Note:** Statistics for parameter estimates reflect the mean and standard deviation of the N=10000 posterior samples of group-level parameters.

If attribute representations in the VMPFC contribute to evidence accumulation, shifting to represent preferences of the choice recipient, what drives changes in those attribute representations, particularly during choices for the *Dissimilar* other? Based on previous research implicating TPJ and STS in directing VMPFC value integration during social decision (e.g., Hare et al., 2010), we speculated that the ERP response for *Dissimilar* choices from 350-400 ms post-stimulus may play a role in perspective taking and attribute weighting. If so, the strength of this response across individuals should predict individual differences in alteration of the weights on taste and/or health. To test this hypothesis, we computed the mean differences in the amplitude of this response for *Dissimilar – Own* and *Similar – Own*, and correlated this difference with differences in taste and health weights for the same contrast. Consistent with the idea that this response may serve to inhibit the use of egocentric preferences, larger ERP amplitude from 350-400 ms across participants significantly predicted reduced weighting on taste for the *Dissimilar* recipient (Spearman’s rho = −.41, P = .01). No other correlations were significant, suggesting a specific role for this signal in reducing egocentric use of taste preferences.

We next sought to use our computational model to understand the nature of the response-locked signals localized to the IPS, which showed the largest difference in signal for *Similar* others (Figure 6D). Notably, applying our computational model to the behavioral data suggested that, rather than resulting from serial correction mechanisms, longer RTs on average when choosing for others could result in part from the adoption of different response thresholds relative to the self, particularly in the case of the *Similar* other (Figure 6E). If so, we would predict that response-locked ERPs localized to the IPS may contribute to adjusting the *response threshold*, rather than correcting egocentric value signals.

Therefore, we directly tested whether this neural response-locked component influenced the threshold parameter by entering the trial-by-trial ERP data from the response-locked posterior SOIs and −300 to −180 ms time window of interest into the M2 specification for the DDM, interacted with the response threshold. Results confirmed a significant relationship between neural response in the IPS and the height of the response threshold (Figure 6F). Moreover, adding this regressor improved model fit (DIC = 41421.1, compared to DIC = 41625.6 for an equivalent to M2 with the same set of trials used in the response-locked ERP version of M2). Importantly, the estimate of the ERP regressor was strongly positive (*P* = .000), with no significant interaction as a function of recipient. Although the polarity of the observed ERP signal depends on a variety of factors including reference electrode, sensor location, and cortical anatomy, the positive influence of this ERP component on the barrier coefficient is in line with the more positive amplitude observed in the response-locked data when choosing for the *Similar* other compared to the other two conditions.

### Simulation exercise: RTs for same and different choices

Above, we demonstrate that the best-fitting parameters of a simple weighted-integration, constant-value DDM can account for overall choice and RT patterns, and can be linked to specific ERP components. Notably, this model supports the idea of egocentric biases, but does *not* include a late-acting mechanism for adjusting preferences when choice for self and similar or dissimilar others diverge. This contrasts with dual-process models such as anchoring-and-adjustment, which are primarily supported by behavioral and RT data. We thus asked whether our model could generate the RT patterns typically associated with an anchoring-and-adjustment mechanism: namely, longer RTs when choices for Similar others diverge from choices for oneself. To do this, we generated 160,000 simulated food choices for each recipient (*Own*, *Similar, Dissimilar*) based on the 16 possible combinations of health and taste ratings. Then, for each pairing of recipients (i.e., *Own* vs. *Similar, Own* vs. *Dissimilar, Similar* vs. *Dissimilar*), we determined how often foods with each combination of taste and health rating led to similar or different simulated choices (Figure 7), as well associated RTs. All statistical comparisons of the simulated RTs were performed using Bayesian analysis incorporating MCMC estimation to generate distributions of likely means and standard deviations (Kruschke, 2013). A P-value of zero thus represents completely non-overlapping simulated distributions.

**Figure 7.**
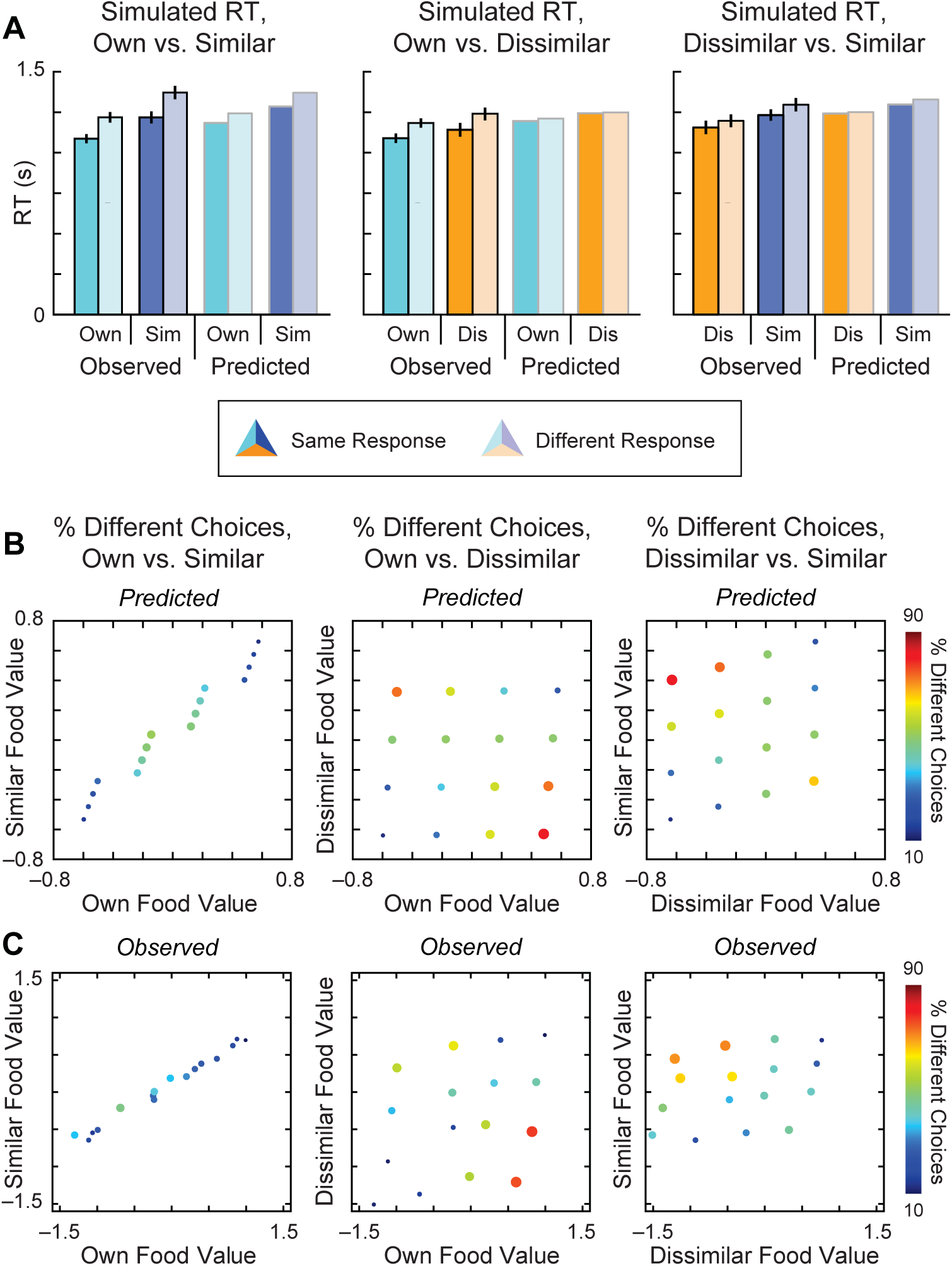
DDM simulations. (A) Simulations (N=10000) of RT data for same versus different choice using best-fitting specification M2. Trials are separated based on the same criteria described for observed data in Figure 2, with observed data (left) plotted next to predicted model RTs (right) for each combination of recipient. (B-C) Predicted (B) and observed (C) relationship between drift rates and percentage of different choices when comparing choices for two recipients, simulated separately for each of the 16 possible combinations of health and taste rating. Proximity of the dots to the diagonal reflects similarity of food values for two recipients, and size and color of dots are coded according to the percentage of different choices. Additional details are provided in the main text.

As expected, model simulations with the fitted parameters suggested that choices for *Own* self and *Similar* other agree more frequently (68% of simulated choices) than for *Dissimilar* other (52% agreement). More importantly, the simulated data also clearly replicate the RT patterns observed both in previous work and in the present study (Figure 7A, top). When comparing decisions for the *Similar* recipient to one’s *Own* choices, RTs were considerably faster for same versus different choices (*P* = .000). When comparing decision for *Dissimilar* other to one’s *Own* choices, RTs were also on average faster for same versus different choices, although the differences were notably smaller (*P* = .073). Finally, when comparing choices for one’s *Own* self to decisions for others, we replicated the novel observation that decisions for the self that result in the same response as that made for similar others *also* tend to be made faster than choices for the self that differ (P = .000).

To provide additional insight into *when* choices converge or diverge, and why these are consistently associated with different RTs, we compared the projected value of a food for two given recipients to the likelihood that the model simulations make different choices about that food (Figure 7B). We divided foods by the sixteen possible combinations of health and taste ratings (e.g., health = 3 and taste = 2). Using health and taste weights derived from model M2, we mapped the projected value of the food versus the percentage of different choices. As expected, this analysis shows that the values of different foods fall closer to the diagonal when comparing choices for one’s *Own* self to a *Similar* other, suggesting that these values are more tightly related (Figure 7B, left). In this case, choices diverge more frequently for options with intermediate overall values (i.e., ratings of Weak No or Weak Yes on both taste and health attributes vs. Strong No or Strong Yes), largely due to noisier choices in the face of indifference. In contrast, when comparing choices made for *Own* self and a *Dissimilar* other (Figure 7B, middle), the values of the food are less correlated, and a greater percentage of divergent choices are observed at the extremes (e.g., very unhealthy but very tasty, or vice versa), due to the differences in attribute weights. Divergent choices in this case are also less likely to be errors.

Next we examined the extent to which observed patterns of choice divergence correspond to our model predictions (Figure 7C). We first used multiple regression to estimate the coefficients on health and taste for choices made in the Own, *Similar*, and *Dissimilar* conditions, separately for each subject. From these coefficients, we estimated the overall decision value for each subject of each food in each condition. Then, for each of the 16 combinations of health and taste ratings, we calculated the average decision value across all subjects for that combination, separately for each condition. Finally, we also calculated the percentage of the time that choices diverged for two recipients, separately for each of the 16 combinations. As can be seen from Figure 7C, the data hew closely to the simulated predictions. Specifically, there is a strong correlation between the value for one’s *Own* self and the value for *Similar* other, and the largest difference between choices for *Own* and *Similar* occur for values close to 0 (Figure 7C, left). In contrast, when comparing values and choices for one’s *Own* self and the *Dissimilar* other, the most divergent choices occur at the extremes of conflict between health and taste (Figure 7C, middle). Comparison of choices for *Similar* and *Dissimilar* lie in between (Figure 7C, right).

Thus, our model explains the longer RTs we observed when making different choices for similar others, compared to dissimilar others. Put simply, in this paradigm people choose differently for similar others in the same cases where they would be less certain what to choose for *themselves*. Our simulations suggest that the extra deliberation time and divergent choice in these trials emerges organically from increased uncertainty within a single decision process, despite giving the appearance of resulting from a serial, slow-to-activate corrective process.

## Discussion

To what extent do decisions for oneself and others rely on shared neurocomputational mechanisms versus recruiting additional slow or deliberative mechanisms (Epley et al., 2004; Tamir and Mitchell, 2013)? Do increases in response time for others reflect effortful correction or uncertainty within a process of stochastic evidence accumulation? Here we exploited the high temporal precision of ERP and the power of computational model fitting (Ruff and Fehr, 2014; Crockett, 2016) to generate novel insights into these important questions.

Consistent with previous work (Nicolle et al., 2012; Janowski et al., 2013), our results strongly support the existence of a shared value integration process for self and others, and further suggest that this process follows similar temporal dynamics. Neural value signals for other recipients emerged within the same temporal window as those for oneself, approximately 500-650 ms after stimulus onset and localized to regions including VMPFC. Importantly, however, these value signals represented taste and health attributes differently depending on the decision context, in line with previous data on dietary self-control (Harris et al., 2013). Moreover, we demonstrated a link between trial-level fluctuations in the ERP value signal and trial-by-trial differences in the process of evidence accumulation associated with each recipient. Whereas VMPFC signals promoted taste considerations when deciding for oneself or the similar other, these same signals promoted health attributes when deciding for the dissimilar, healthy eater. Notably, these signals emerged in the stimulus-locked data more than 400 ms prior to the average response time, suggesting that they represent an input into brain systems associated with choice selection (Hare et al., 2011) rather than directly implementing accumulation-to-bound and choice at the motor level. However, in light of previous work linking VMPFC response to choice selection processes (Hunt et al., 2012; Strait et al., 2014), as well as other data demonstrating rapid interactions between VMPFC and sensorimotor systems (Harris and Lim, 2016), further research will be necessary to fully disentangle the relationship between ERP value signals and evidence accumulation.

This shared VMPFC mechanism might also explain, in part, the robust and consistent evidence for egocentric biases when choosing for others (Ross et al., 1977; Devaine and Daunizeau, 2017; Tarantola et al., 2017). In our study, we found some evidence that people use their own preferences to guide choices for others, particularly when those preferences may be relevant. One’s own weights on *both* taste and health predicted weights used when choosing for a similar other, whereas when choosing for a dissimilar, healthy eater, only one’s own health preferences predicted choice. Thus, we find not only that people use their own preferences, but also that they do so selectively based on inferences about the intended recipient.

Our results also support the use of additional perspective-taking mechanisms, particularly when deciding for people with different preferences from one’s own. We observed an ERP component specific to choosing for the *Dissimilar* recipient from 350 to 400 ms after stimulus onset, just before the time window associated with stimulus valuation. This signal was localized to posterior temporal areas near the temporoparietal junction (TPJ), a region previously implicated in various aspects of social cognition, including suppression of egocentric biases (Silani et al., 2013), perspective taking (Saxe, 2006; Tusche et al., 2016), and pro-social behavior (Morishima et al., 2012; Hutcherson et al., 2015). However, the timing of this response, as well as the failure of time-varying egocentrically-biased computational models to better account for choice and RT patterns, are both inconsistent with the idea that such perspective-taking mechanisms activated slowly, *after* the emergence of a default self-centered bias (Epley et al., 2004; Tamir and Mitchell, 2013).

Nonetheless, we do find evidence for late-acting, response-related control mechanisms. We observed a comparatively late ERP signal differentiating choices by recipient from −300 ms before response. Whereas the stimulus-locked component showed an increased response specifically for the *Dissimilar* recipient, this neural activity predominantly distinguished the *Similar* condition. Yet, in contrast to the idea that this signal might be needed to correct or inhibit egocentric values, our computational analyses linked it only to variation in the response threshold for choice. What could be driving increases in response threshold for the similar recipient? One possible factor could be choice uncertainty. In line with this idea, participants took longer on average and had higher decision thresholds when choosing for the similar other, suggesting the need for additional evidence accumulation relative to the other conditions. Additionally, weights on taste were modestly but consistently lower for the similar other compared to choices for one’s self, while weights on health were slightly higher, suggesting that participants may have “hedged” their choices where the similar other was concerned. These results provide a different perspective on the role of controlled processing in self-other decision-making. They suggest that some forms of controlled processing may influence choices for others not by modifying value signals, but by raising the criterion for choice.

Given evidence of strong egocentric biases, rapid perspective-taking, and no computational evidence for delayed adjustment, what then explains patterns of longer response time when choosing differently from the self? Simulations performed using our combined neurocomputational model suggested that choices for self and other can diverge either due to differences in attribute weighting (as observed when making choices for the dissimilar other), or simply due to noise in the process of evidence accumulation (as observed when making choices for partners with similar preferences). In other words, trials where choices for self and similar others diverge may sometimes be trials on which choices for the self alone might be inconsistent from one instantiation to the next, due to ambivalent and noisy values. These will also tend to be the choices with the longest RTs, for both self and others. Although our model underestimated the observed RT differences somewhat, we suspect that the extra time may stem from conflict-triggered increases in threshold at the single-trial level (Cavanagh et al., 2011). This effect would tend to magnify the difference in RT beyond that predicted by a canonical DDM, in precisely those trials where divergent choices are most likely to be observed. Taken together, these results suggest that, at least in some decision scenarios, patterns of behavior previously associated with the action of both anchoring and adjustment may be better explained simply by anchoring, implemented in a stochastic decision system.

Note that these results do not argue against the basic idea of dual- or even multi-system models of choice. If anything, our results highlight the importance of *both* perspective-taking *and* response inhibition, and tie them to value integration and threshold computations, respectively. Moreover, there may be important decision scenarios in which choices for others do engage dissociable correction or perspective-taking mechanisms, operating at different timescales (Epley et al., 2004; Evans, 2008). For example, our paradigm included no feedback. In cases where a decision maker must integrate both general information (e.g., knowledge about a person’s culture, background, or similarity to oneself) and direct experience with another’s choices, we might expect an initial egocentric preference that is corrected based on learning and memory (Devaine and Daunizeau, 2017; Tarantola et al., 2017). There may also be important individual differences in the speed or strength of social cognitive mechanisms, which might depend on the integrity of areas like the STS and TPJ. Our work highlights the utility of computational modeling for determining when patterns of choice, RT, and neural data that appear on their face to support dual-process models are instead more consistent with a single value integration mechanism (as observed in our study) and when additional computational mechanisms might be necessary to fully account for behavior.

Careful consideration of the timing of these processes also generates a novel prediction regarding the effects of time pressure versus cognitive load in social decision scenarios. Our data suggest that making accurate choices for people with divergent preferences relies more on early perspective-taking mechanisms, occurring from 300 to 500 ms post-stimulus onset in TPJ, while making accurate choices for partners with uncertain preferences relies more on later response threshold mechanisms localized to posterior cingulate cortex. If this is indeed the case, manipulations using time pressure or cognitive load may sometimes have quite different effects on social decision-making. Time pressure, which in perceptual choice has been tied to reductions in response threshold (Forstmann et al., 2010), might be expected to result in reduced accuracy only when others’ preferences are uncertain. In contrast, cognitive load may disrupt effortful perspective-taking mechanisms, resulting in stronger reductions in accuracy when preferences differ from the self.

Our work provides the first computational account of decision making capable of explaining the full pattern of choice, response times and neural dynamics when choosing for others, in a context where preferences are well-specified but can differ among individuals. Future studies will need to examine choices for others in different contexts and with different information. We believe that the unique combination of neural and computational modeling presented here provides a powerful and essential tool for this research, and will foster exciting new avenues for exploring how we choose for ourselves and others.

## Author Contributions

A.H. and C.A.H. designed the experiment. A.H. collected the data. A.H., J.A.C., and C.A.H. analyzed the data. A.H., J.A.C., and C.A.H. interpreted the results and wrote the paper.

